# Hedgehog signaling activates a heterochronic gene regulatory network to control differentiation timing across lineages

**DOI:** 10.1101/270751

**Authors:** Megan Rowton, Carlos Perez-Cervantes, Ariel Rydeen, Suzy Hur, Jessica Jacobs-Li, Nikita Deng, Emery Lu, Alexander Guzzetta, Jeffrey D. Steimle, Andrew Hoffmann, Sonja Lazarevic, Xinan Holly Yang, Chul Kim, Shuhan Yu, Heather Eckart, Sabrina Iddir, Mervenaz Koska, Erika Hanson, Sunny Sun-Kin Chan, Daniel J. Garry, Michael Kyba, Anindita Basu, Kohta Ikegami, Sebastian Pott, Ivan P. Moskowitz

## Abstract

Heterochrony, defined as differences in the timing of developmental processes, impacts organ development, homeostasis, and regeneration. The molecular basis of heterochrony in mammalian tissues is poorly understood. We report that Hedgehog signaling activates a heterochronic pathway that controls differentiation timing in multiple lineages. A differentiation trajectory from second heart field cardiac progenitors to first heart field cardiomyocytes was identified by single-cell transcriptional profiling in mouse embryos. A survey of developmental signaling pathways revealed specific enrichment for Hedgehog signaling targets in cardiac progenitors. Removal of Hh signaling caused loss of progenitor and precocious cardiomyocyte differentiation gene expression in the second heart field *in vivo*. Introduction of active Hh signaling to mESC-derived progenitors, modelled by transient expression of the Hh-dependent transcription factor GLI1, delayed differentiation in cardiac and neural lineages *in vitro*. A shared GLI1-dependent network in both cardiac and neural progenitors was enriched with FOX family transcription factors. FOXF1, a GLI1 target, was sufficient to delay onset of the cardiomyocyte differentiation program in progenitors, by epigenetic repression of cardiomyocyte-specific enhancers. Removal of active Hh signaling or *Foxf1* expression from second heart field progenitors caused precocious cardiac differentiation *in vivo*, establishing a mechanism for resultant Congenital Heart Disease. Together, these studies suggest that Hedgehog signaling directly activates a gene regulatory network that functions as a heterochronic switch to control differentiation timing across developmental lineages.

## INTRODUCTION

Developmental processes are governed by embryonic timing mechanisms and tight control over differentiation timing is critical for organ development (Ebisuya and Briscoe, 2018). Developmental timing affects cell type identity (Kohwi and Doe, 2013; Pearson and Doe, 2004), spatial patterning (Gomez et al., 2008; Harima et al., 2013; Hubaud and Pourquié, 2014; Schröter et al., 2008), proliferation rate (Momen-Roknabadi et al., 2016; Pourquié, 1998) and differentiation rate (Brown, 2014; Imayoshi et al., 2013; Moss, 2007; Otani et al., 2016; Weger et al., 2017). While the concept of heterochrony - differences in the timing or duration of developmental processes - was developed to explain developmental distinctions between species, it has also provided a theoretical framework for researchers investigating the etiologies of congenital malformations in humans and animal models (Smith, 2003; Wilson, 1988). An appreciation of the intrinsic and extrinsic regulatory mechanisms controlling the timing of developmental transitions is crucial to understanding organ development, organ homeostasis and the etiology of developmental disorders.

Initial evidence for the explicit genetic control of developmental timing was first provided in *C. elegans*, through the identification of heterochronic mutations that caused either premature or delayed developmental transitions (Ambros and Horvitz, 1984; reviewed in Moss, 2007). Mutations in the *C. elegans* gene *lin-4*, for example, caused a reiteration of larval stages, thereby delaying differentiation (Ambros and Horvitz, 1984). Conversely, mutations in the genes *lin-28* and *lin-14* caused premature differentiation of progenitor cells (Ambros, 1989; Ambros and Horvitz, 1984; Euling and Ambros, 1996; Harandi and Ambros, 2015; Moss et al., 1997; Ruvkun and Giusto, 1989). These and other studies exploited the invariant cell lineages of *C. elegans* to identify a class of heterochronic mutations that specifically disrupted the temporal execution of differentiation, uncoupling control of differentiation rate from specification or patterning of cell lineages. This characteristic is a defining feature of heterochronic regulatory networks (Smith, 2003).

Developmental timers linked to specification or patterning have been identified in some contexts, such as the *Drosophila* neuroblast temporal identity factors (Averbukh et al., 2018; Durand and Raff, 2000; Jessell TM, 2000; Pearson and Doe, 2004) or the oscillatory network controlling vertebrate somitogenesis (Pourquié, 1998). However, heterochronic modulators of differentiation rate have proven difficult to identify in complex vertebrate embryos (Johnson and Day, 2000). Mammalian orthologues of the *C. elegans* heterochronic gene *lin-28* maintain stem cell properties and delay terminal differentiation, suggesting the possibility of a homologous role (Balzer et al., 2010; Komarovsky Gulman et al., 2019; Shyh-Chang and Daley, 2013; Thornton and Gregory, 2012; Urbach et al., 2014; Viswanathan and Daley, 2010; West et al., 2009; Yermalovich et al., 2019; Yu et al., 2007; Yuan et al., 2012; Zhang et al., 2016). Despite an abundance of model systems displaying species-specific differences in differentiation timing, however, few other candidates for mammalian heterochronic regulation have been identified, particularly in lineage-specified progenitor populations (Van den Ameele et al., 2014; La Manno et al., 2016; Otani et al., 2016). Furthermore, while known examples implicate cell autonomous transcriptional modules in the regulation of differentiation timing, the highly-coordinated differentiation of mammalian organs, such as the heart and lungs (Peng et al., 2013; Steimle et al., 2018) suggests that intercellular signaling pathways may serve a heterochronic function.

The vertebrate heart forms from two pools of cardiac progenitors (CPs), the first heart field (FHF) and the second heart field (SHF), defined by their differentiation order. The FHF generates the linear heart tube (HT), a functional pump comprised of differentiating cardiomyocytes (CMs) reminiscent of the rudimentary heart observed in invertebrates. The SHF remains a reservoir of undifferentiated CPs that subsequently migrate into the HT and differentiate later (Kelly, 2012). SHF progenitors generate the right ventricle, pulmonary outflow tract, and atrial septum, cardiac structures critical for cardiopulmonary circulation and commonly affected in Congenital Heart Disease (CHD) (Briggs et al., 2012; Kelly et al., 2014; Neeb et al., 2013). Both the FHF and the SHF appear to be specified toward the cardiac lineage in the early embryo (E7.25 in the mouse embryo) (Dyer and Kirby, 2009; Lescroart et al., 2014; Meilhac et al., 2004), however, the molecular basis of the delayed differentiation of the SHF relative to the FHF remains unclear. SHF progenitors are exposed to a complex intersection of signaling pathways that promote various progenitor behaviors and disruption of SHF signaling pathways often causes CHD (Hutson et al., 2010; Jain et al., 2015; Peng et al., 2013; Rochais et al., 2009; Sorrell and Waxman, 2011). We investigated the hypothesis that intercellular signaling pathways provide heterochronic control of CP differentiation timing in the SHF that is critical for cardiac morphogenesis.

We report that Hedgehog (Hh) signaling drives a heterochronic gene regulatory network (GRN) controlling the rate of mammalian CM differentiation. SHF CPs demonstrate selective Hh signaling activity that is lost in differentiating CMs *in vivo* and *in vitro*. Inactivation of Hh signaling in the SHF results in loss of progenitor gene expression and premature activation of the cardiac differentiation program. Maintained expression of the activating Hh pathway transcription factor (TF) GLI1 in CPs or neural progenitors delays their differentiation. Hh signaling activates a progenitor-specific GRN enriched in FOX family TFs in CPs, neural progenitors, and in a medulloblastoma model, suggesting that Hh signaling may be broadly deployed to delay differentiation across germ layers and biological contexts. We uncover a Hh-dependent heterochronic molecular mechanism in CPs, in which the GLI1 target FOXF1 is sufficient to directly repress cardiac gene expression by binding to, and epigenetically repressing, CM-specific regulatory elements. Removal of Hh signaling or *Foxf1* in the SHF causes premature cardiac differentiation and CHD *in vivo*, providing a mechanism for CHD. These findings identify the Hh signaling pathway as a cell non-autonomous heterochronic regulator of mammalian differentiation timing.

## RESULTS

### A Screen of Intercellular Signaling Pathway Activity During Cardiomyocyte Differentiation Using Single Cell Transcriptional Profiling

In an effort to define signal-dependent mechanisms that control cardiac differentiation timing, we investigated the activity of candidate developmental signaling pathways across a trajectory of CM differentiation. We analyzed single-cell transcriptomes using droplet-based single-cell RNA sequencing (Drop-seq) (Macosko et al., 2015) on microdissected SHF and heart inflow tract (HT) regions from wild type E10.5 embryos (Figure A). Clustering of the transcriptomes of 3,824 cells from two biological replicates defined 19 distinct populations expressing markers of known cell types from all three germ layers (Figure 1B and Supplemental Table 1). Cluster-wise distribution of cells, and number of detected unique molecular identifiers (UMIs) and genes were well-correlated between replicates (Supplemental Figure 1A-C). With iterative clustering of sub-populations, we focused on four cardiac-related clusters, including two posterior SHF (pSHF) CP populations, an intermediate CP/CM population and a CM population based on cluster-specific differential gene expression patterns (Supplemental Figure 1D-H, see Methods). A differentiation continuum of single cell transcriptomes from CPs in the pSHF to functionally differentiated HT CMs in the heart was apparent (Figure 1C and Supplemental Figure 2A). The two CP clusters expressed known pSHF genes *Isl1, Tbx5* and *Arg1* (Figure 1D) (Cai et al., 2003; de Soysa et al., 2019; Xie et al., 2012) and represented right (CP1) and left (CP2) pSHF populations based on differential expression of the right-sided marker *D030025E07Rik/Plyrr* lncRNA (Welsh et al., 2015) and left-sided marker *Pitx2* (Chengyu Liu, 2002; K. Kitamura, 1999) (Supplemental Figure 2B-D and Supplemental Table 2). The intermediate CP/CM cluster (intermediate) demonstrated high expression of the cardiogenic transcription factor kernel *Tbx5, Gata4* and *Nkx2-5* and low expression of genes encoding sarcomere structural components including cardiac troponin T2 (*Tnnt2*), and the differentiated CM cluster (CM) showed high expression of the cardiogenic kernel and *Tnnt2*.

**Figure 1.**
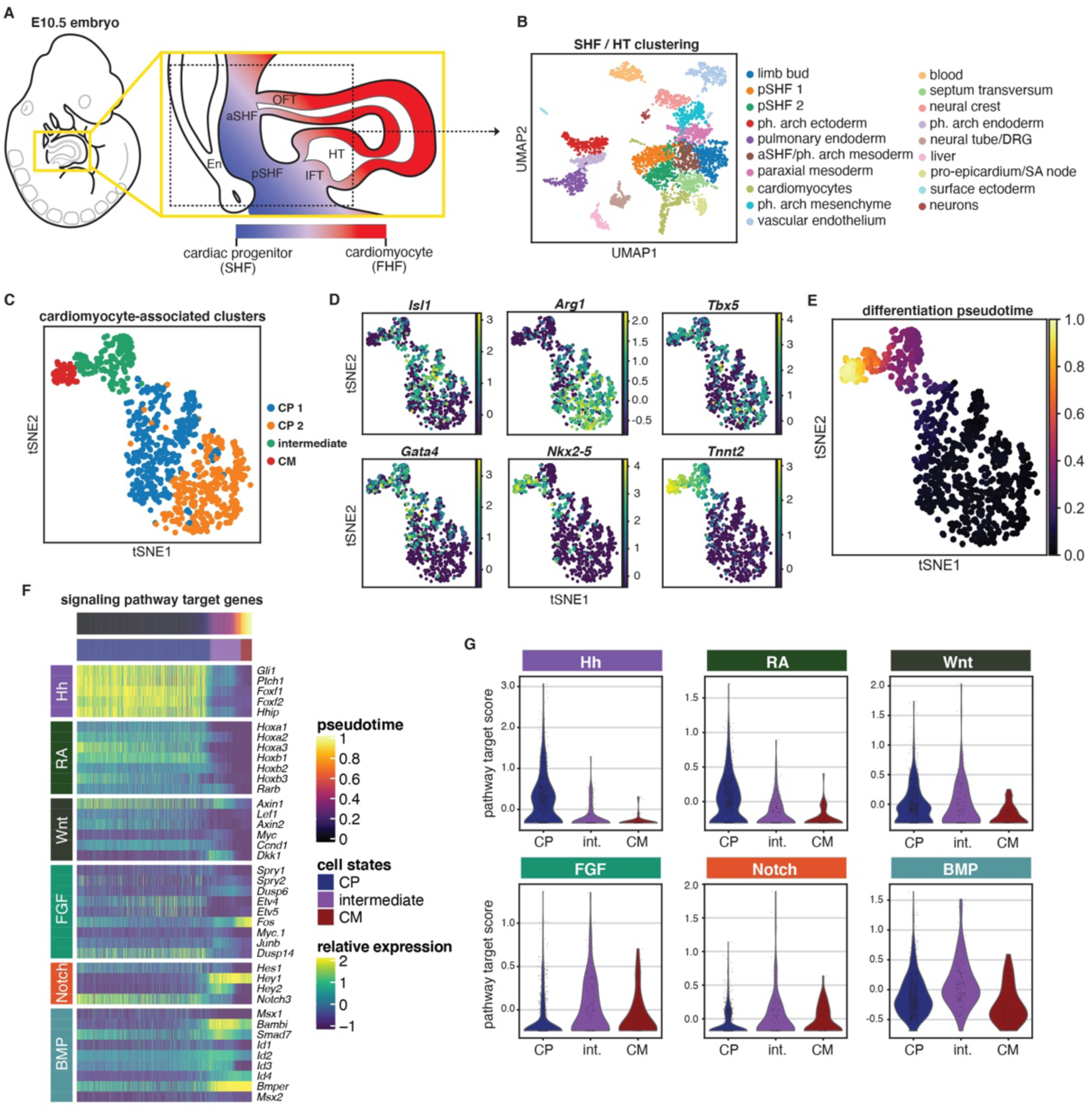
Hedgehog signaling is highly active and specific to the cardiac progenitors of the embryonic second heart field. **(A)** Diagram of the SHF and HT regions of an E10.5 mouse embryo. Dotted lines indicate the boundaries of microdissection for Drop-seq. aSHF, anterior second heart field; pSHF, posterior second heart field; OFT, outflow tract; IFT, inflow tract; HT, heart tube, En, pulmonary endoderm. **(B)** UMAP plot displaying 19 distinct cell type clusters identified from microdissected SHF and HT tissue. **(C)** tSNE plot showing 4 CM-associated clusters of pSHF and HT cells. CP, cardiac progenitor; CM, cardiomyocyte. **(D)** tSNE plots indicating the expression levels of marker genes of CPs (*Isl1, Tbx5, Arg1*), intermediate cells (*Tbx5, Nkx2-5, Gata4*) and CMs (*Tbx5, Nkx2-5, Gata4, Tnnt2*) within the 4 CM-associated clusters. **(E)** tSNE plot of 4 CM-associated clusters with cells colored based on cardiac differentiation pseudotime (dpt) score. Values shown for each cell are calculated as 1-dpt. **(F)** Heatmap of individual cells from CM-associated Drop-seq clusters showing the denoised expression levels of target genes of 6 signaling pathways along the pseudotime differentiation trajectory. Cells are ordered on the x-axis by pseudotime and are clustered on the y-axis by signaling pathway. CP, cardiac progenitor; CM, cardiomyocyte. **(G)** Violin plots showing the metagene scores for signaling pathway target genes at the CP, intermediate and CM stages of cardiac differentiation for 6 signaling pathways. CP, cardiac progenitor; int., intermediate; CM, cardiomyocyte.

We defined a CM differentiation trajectory from CP to intermediate to CM populations from a differentiation pseudotime score for each cell from the four CM-associated clusters. Using a set of 24 known CM marker genes (Supplemental Table 3) (de Soysa et al., 2019) to perform a metagene analysis, we calculated a “cardiac differentiation score” for each cell within the 4 CM-associated clusters. We assigned the cell with the highest cardiac differentiation score as the starting point for a reverse-differentiation pseudotime analysis and observed a smooth trajectory from the combined CP clusters to the intermediate cluster to the CM cluster (Figure 1E and Supplemental Figure 2E). This trajectory was then confirmed by cell-cycle and gene expression analysis. First, we observed a step-wise decrease in cell cycle progression from CP to intermediate to CM stages, consistent with a gradient of increasing differentiation at the expense of proliferation (Supplemental Figure 2F-G). Next, we confirmed expected gene expression differences between the CP and CM clusters by performing a pairwise differential expression test. We found 3,524 CM-enriched and 4,543 CP-enriched genes (*p-*value < 0.05) (Supplemental Table 4). CM-enriched genes such as *Tnnc1, Myl4, Nppa* and *Des* were highly expressed in cells at the end of the pseudotime differentiation trajectory and were associated with GO terms consistent with mature CMs, such as muscle structure development and actin binding (Supplemental Figure 2H-I). In contrast, CP-enriched genes *Isl1, Pdgfra* and *Ccnd1* were more highly expressed in cells at the beginning of the pseudotime trajectory, and associated with GO terms consistent with a progenitor state, including cell division and chromatin organization (Supplemental Figure 2J-K). Furthermore, analysis of the expression patterns of known CP and CM genes along the pseudotime axis supported the pseudotime differentiation trajectory (Supplemental Figure 2L-M) (Bax et al., 2009, 2010; Bertrand et al., 2011; Fujii et al., 2017; Lu et al., 1998; Robb et al., 1998; de Soysa et al., 2019). These observations confirm that the differentiation pseudotime trajectory accurately delineates a cardiomyocyte differentiation pathway.

We attempted to identify signaling pathways with progenitor specific activity in CPs. We investigated developmental signaling pathways known to be active in the SHF, including the Wnt, FGF, Hh, Notch, retinoic acid (RA) and BMP pathways, for local enrichment along the CM differentiation pseudotime trajectory, using expression levels of known target genes from each pathway as a readout of pathway activity (Bruneau, 2013; Han et al., 2019; Hutson et al., 2010; Jain et al., 2015; Rochais et al., 2009; van Wijk et al., 2007). We found that Hh, RA and Wnt pathway targets were active in the CP state at the beginning of the pseudotime trajectory, FGF and Notch pathway targets became activated in the late CP-state and intermediate states, and BMP targets became active in the intermediate and CM states at the end of the pseudotime trajectory (Figure 1F). We observed that Hh pathway targets were most highly expressed in the CP state and were also the most CP-specific targets when we aggregated the expression levels of pathway target genes (Figure 1G and Supplemental Figure 2N). The abrupt loss of Hh signaling activity during the transition of cells from the CP stage to the intermediate stage suggested a specific role for Hh signaling in CPs, and is consistent with a known requirement for Hh signaling in the mammalian SHF during cardiac morphogenesis (Briggs et al., 2016; Goddeeris et al., 2007; Hoffmann et al., 2009; Xie et al., 2012).

### Hh-signaling Promotes CP and Inhibits CM Gene Expression in the pSHF

The molecular consequences of Hh signaling in pSHF CPs, received non-cell autonomously from the adjacent pulmonary endoderm (Goddeeris et al., 2008; Hoffmann et al., 2009), have not been thoroughly described. To uncover the *Shh*-dependent network deployed in CPs, we performed transcriptional profiling of the microdissected pSHF from *Shh* mutant (*Shh*^-/-^) and control (*Shh*^+/+^) littermate embryos at E10.5 (Figure 2A and Supplemental Figure 3A-B). We observed repression of mesoderm- and CP-specific genes such as *Foxf1*, *Wnt2b*, *Snai1,* and *Osr1* in the *Shh*^-/-^ pSHF by RNA-seq (log_2_ FC ≥ 0.5, FDR ≤ 0.05) and confirmed by qPCR or *in situ* hybridization (Figure 2B, Supplemental Figure 3C-D and Supplemental Table 5). In contrast, we observed activation of genes encoding CM differentiation products in the *Shh*^-/-^ pSHF, including sarcomere components *Myl3*, *Tnni3,* and atrial specific markers such as *Nppa* (Figure 2B and Supplemental Figure 3C). These observations suggested that active Hh signaling in the SHF promoted CP gene expression and inhibited CM gene expression. To test this hypothesis more definitively, we compared genes exhibiting Hh-dependence in the *Shh^-/-^* pSHF with all genes exhibiting CP- or CM-enriched gene expression, defined with transcriptional profiling of the wild type pSHF versus HT at E10.0 (log_2_ FC ≥ 0.5, FDR ≤ 0.05) (Supplemental Table 6). This analysis revealed that Hh-activated genes, repressed in the *Shh*^-/-^ SHF, were preferentially expressed in the wild type SHF compared to the wild type HT (*p* = 2e^−16^) (Figure 2C). Conversely, Hh-repressed genes, activated in the *Shh*^-/-^ SHF, were preferentially expressed in the wild type HT compared with the wild type SHF (*p* = 2e^−16^). These results confirmed that Hh signaling promoted pSHF-specific gene expression and repressed HT-specific gene expression in the pSHF.

**Figure 2.**
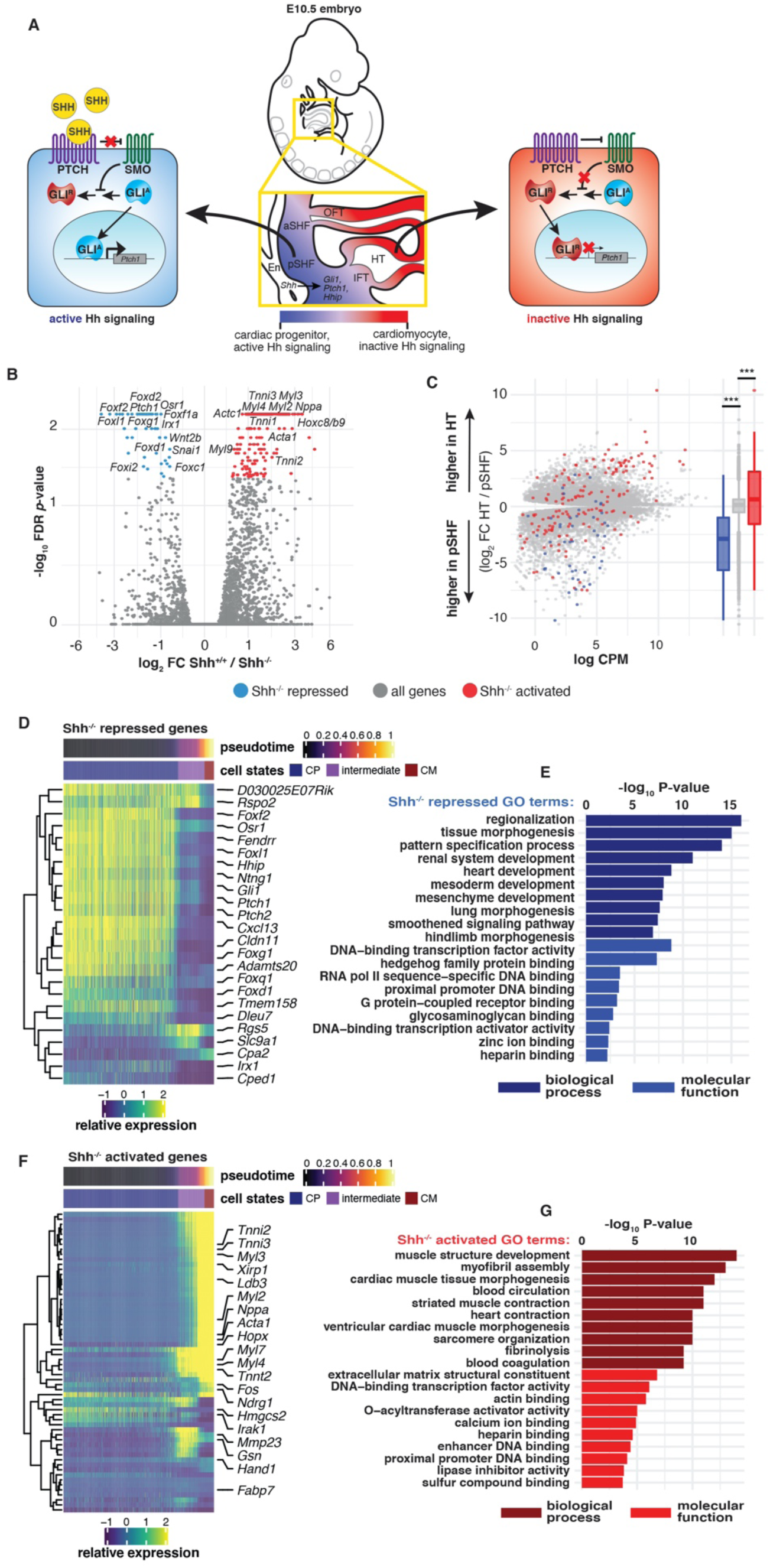
Hedgehog signaling activates a progenitor gene program and represses the cardiac differentiation program, in the SHF. **(A)** Model of Hh signaling in CPs (blue) in the pSHF and differentiating CMs (red) in the HT from the E10.5 mouse embryo. Active Hh signaling in CPs causes accumulation of GLI^A^ in the nucleus and the activation of Hh target genes, such as *Ptch1*, in CPs (left). Inactive Hh signaling results in the truncation of GLI^A^ to GLI^R^ and accumulation of GLI^R^ repressor in the nuclei of CMs (right). **(B)** Volcano plot displaying activated and repressed genes in the *Shh^-/-^* pSHF relative to *Shh^+/+^* embryos. Red and blue dots signify significantly dysregulated genes (log_2_ FC > 0.6, FDR < 0.05). **(C)** MA plot and box plots illustrating the distribution of *Shh^-/-^* dysregulated genes in the context of differential expression between the wild type pSHF and HT. ANOVA for three groups: *F-value* = 246.2, *p-value* < 2e^−16^; post-hoc pairwise T-test with Benjamini-Hochberg correction: activated vs all genes *p-value* = 1e^−12^, repressed vs all genes *p*-value < 2e^−16^. **(D)** Heatmap showing denoised data from individual cells from cardiac-associated Drop-seq clusters indicating the wild type expression levels of *Shh^-/-^* repressed genes along the pseudotime differentiation trajectory. Cells are ordered on the x-axis by pseudotime and are hierarchically clustered on the y-axis. CP, cardiac progenitor; CM, cardiomyocyte. **(E)** Gene ontology (GO) analysis of *Shh^-/-^* repressed genes. **(F)** Heatmap showing denoised data from individual cells from cardiac-associated Drop-seq clusters indicating the wild type expression levels of *Shh^-/-^* activated genes along the pseudotime differentiation trajectory. Cells are ordered on the x-axis by pseudotime and are hierarchically clustered on the y-axis. CP, cardiac progenitor; CM, cardiomyocyte. **(G)** Gene ontology (GO) analysis of *Shh^-/-^* activated genes.

Hh-dependent promotion of CP and repression of CM gene expression programs predicted a rapid differentiation switch along the pSHF CP to CM differentiation trajectory from the Drop-seq analysis. In fact, we observed that with few exceptions, *Shh^-/-^* repressed genes from bulk RNA-seq were exclusively expressed in the CP stage at the beginning of the pseudotime differentiation trajectory (Figure 2D), and were associated with GO terms such as Smoothened (Hh) signaling and embryonic developmental terms including regionalization, pattern specification and heart development (Figure 2E). In contrast, the majority of genes activated in the *Shh^-/-^* pSHF were expressed only in the intermediate and CM states (Figure 2F), and were associated with GO terms such as muscle cell differentiation, myofibril assembly, sarcomere, myofibrils, and cardiac muscle tissue development (Figure 2G). Together, these findings confirmed that Hh signaling is required to maintain CP-specific gene expression and suppress CM differentiation gene expression in the pSHF, implicating the Hh pathway as a candidate heterochronic regulator of CM differentiation timing.

### Hh Signaling Delays Differentiation of CPs Derived from Mouse Embryonic Stem Cells

We hypothesized that a transition from active Hh signaling to inactive Hh signaling was required to permit CM differentiation. We therefore predicted that transiently exposing CPs to active Hh signaling, as they experience in the SHF, would inhibit CM differentiation. We tested this prediction using an *in vitro* cardiac differentiation system in which mouse embryonic stem cells (mESCs) were directed to undergo CM differentiation via biologically relevant intermediates, including mesoderm (Day 3, D3), CPs (D5), and immature CMs (D8+), expressing known markers of each stage (Supplemental Figure 4A-B) (Kattman et al., 2011). We generated a mESC line with doxycycline-inducible expression of GLI1, an activating GLI TF (GLI^A^) that drives Hh-dependent target gene expression (Ruiz i Altaba et al., 2007), fused to a C-terminal FLAG-TEV-AVI tag (GLI1-FTA), to model the induction of active Hh signaling (Figure 3A). We first evaluated the endogenous activity of Hh signaling in the absence of GLI1 induction during *in vitro* cardiac differentiation of the GLI1-FTA cell line. We observed a decrease in Hh signaling activity as cells transitioned from mesoderm to CP and CM stages during the mESC differentiation time-course. Hh signal transduction is effected by the accumulation of activating GLI TFs (Gli^A^ TFs, e.g. GLI1) that activate a set of context-independent target genes, including *Gli1* itself, *Ptch1* and *Hhip*, which serve as canonical markers of active Hh signaling (Dai et al., 1999; Hui and Angers, 2011; Lee et al., 1997; Ruiz i Altaba et al., 2007). mESCs and mesoderm stage cells exhibited *Gli1* and *Ptch1* transcript expression and abundance of the Gli^A^ TF, GLI1, however, these markers of active Hh signaling were depleted during differentiation towards CPs (D5-7) and differentiated CMs (D8-12) (Supplemental Figure 4C-D). A second metric of active Hh signaling is the accumulation of activating GLI TFs (Gli^A^ TFs, e.g. GLI1) at the expense of truncated, repressive GLI TFs (Gli^R^ TFs, e.g. truncated GLI3 (GLI3^R^)), and the Gli^A^ / Gli^R^ ratio provides a quantitative metric of Hh signaling activity (Ruiz i Altaba et al., 2007). Quantification of the relative protein abundances of GLI3^A^ versus GLI3^R^ isoforms demonstrated a switch from active Hh signaling to inactive Hh signaling coincident with CM differentiation, assessed by western blot (Supplemental Figure 4E-F). These observations indicated that a transition from active to inactive Hh signaling was coincident with CM differentiation *in vitro*.

**Figure 3.**
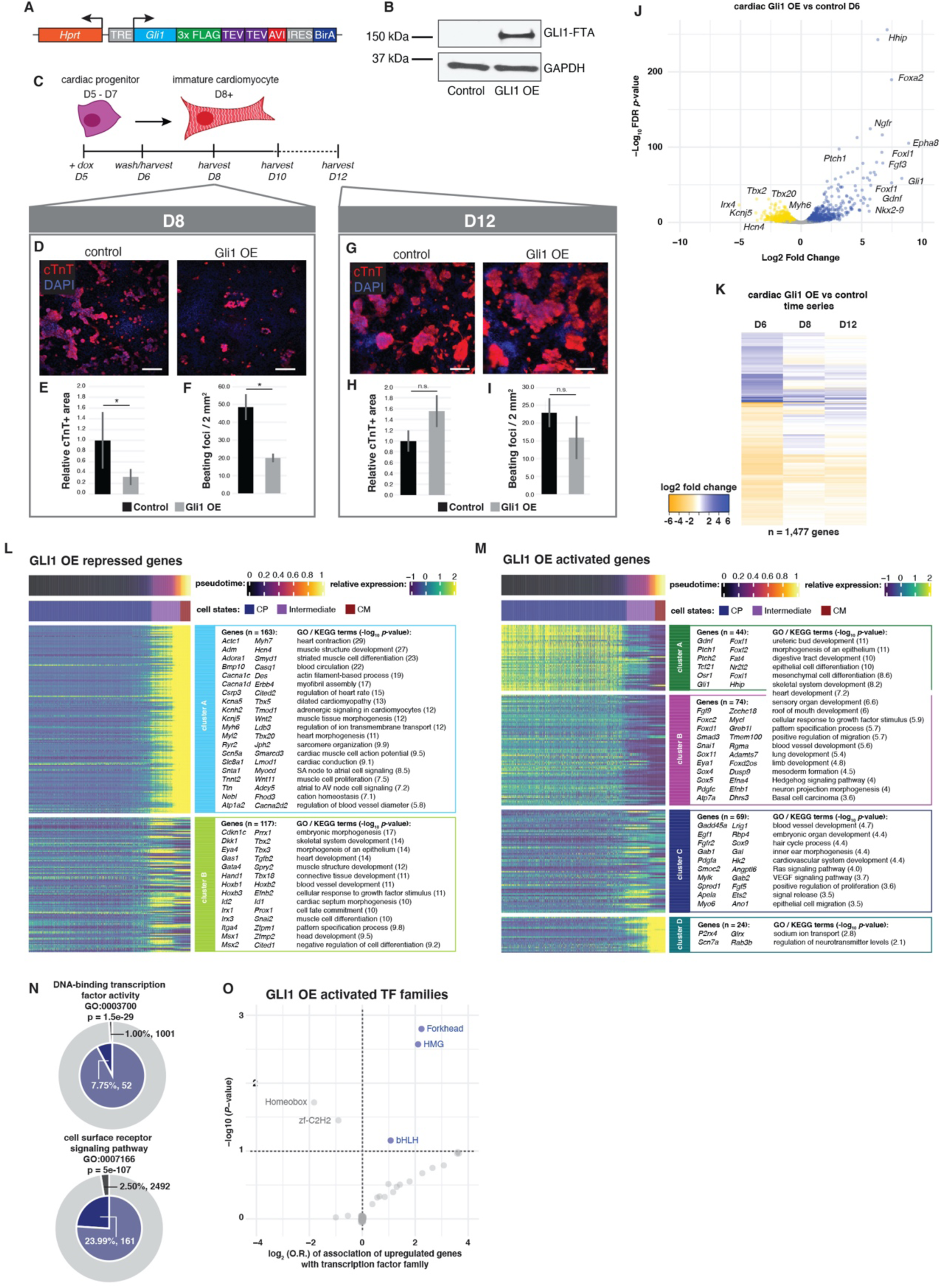
GLI1 expression in CPs delays cardiomyocyte differentiation and activates a progenitor network enriched with FOX TFs. **(A)** Diagram of the doxycycline-inducible GLI1-FTA transgenic cassette inserted into the *Hprt* locus in mESCs. **(B)** Western blot showing induction of GLI1 protein, a marker of active Hh signaling, after doxycycline treatment for 24 hours in CPs. **(C)** Schematic representation of the transient GLI1 OE experimental design employed in mESC-derived differentiating CMs. Cells were treated with doxycycline in the CP state from D5-6, doxycycline was removed at D6, and cells were permitted to continue differentiation until the CM state at D8 or D12. **(D)** Immunofluorescent staining for cardiac troponin (cTnT) in control and GLI1 OE cells harvested at D8. cTnT staining is shown in red and DAPI counterstain is shown in blue. Scale bar = 100μm. **(E)** Quantification of the area of cTnT-positivity in control and GLI1 OE cells at D8 (Students T-test, * *p*-value ≦ 0.05). **(F)** Quantification of the number of beating foci in videos taken of control and GLI1 OE cells at D8. (Students T-test, * *p*-value ≦ 0.05). **(G)** Immunofluorescent staining for cardiac troponin in control and GLI1 OE cells harvested at D12. cTnT staining is shown in red and DAPI counterstain is shown in blue. Scale bar = 100μm. **(H)** Quantification of the area of cTnT-positivity in control and GLI1 OE cells at D12. (Students T-test, n.s. = not significant). **(I)** Quantification of the number of beating foci in videos taken of control and GLI1 OE cells at D12. (Students T-test, n.s. = not significant). **(J)** Volcano plot displaying activated and repressed genes in the GLI1 OE CPs relative to control CPs embryos at D6. Blue and yellow dots signify significantly activated and repressed genes, respectively (log_2_ FC > 0.5, FDR < 0.05). **(K)** Heatmap time-series of the log_2_ FC values of genes differentially expressed in GLI1 CP OE relative to control differentiations at D6, D8 and D12. **(L)** Heatmap of individual cells from cardiac-associated Drop-seq clusters displaying denoised wild type expression levels of GLI1 OE repressed genes along the pseudotime differentiation trajectory. Cells are ordered on the x-axis by pseudotime and are clustered on the y-axis with k-means clustering (k=2). GO terms associated with repressed genes in each cluster are also shown. CP, cardiac progenitor; CM, cardiomyocyte. **(M)** Heatmap of individual cells from cardiac-associated Drop-seq clusters displaying denoised wild type expression levels of GLI1 OE activated genes along the pseudotime differentiation trajectory. Cells are ordered on the x-axis by pseudotime and are clustered on the y-axis using k-means clustering (k=4). GO terms associated with activated genes in each cluster are also shown. CP, cardiac progenitor; CM, cardiomyocyte. **(N)** Nested pie-chart demonstrating enrichment of genes associated with DNA-binding TF activity (*p* = 1.5e^−29^) and cell surface receptor signaling pathways (*p* = 5e^−107^) within GLI1 OE activated genes relative to a background of all expressed genes (TPM >1). **(O)** Dot plot showing enrichment (log_2_ odds ratio (O.R.) vs −log_10_ *p*-value) of genes encoding TFs in the Forkhead (FOX), HMG and bHLH families within genes activated by GLI1 OE.

To investigate whether active Hh signaling delayed CM differentiation, we modeled CP-specific activation of Hh signaling by inducing a pulse of GLI1-FTA expression at the CP stage of the mESC-cardiac differentiation (D5 to D6), mimicking the transient exposure of pSHF CPs to active Hh signaling. Treatment of D5 CPs with 500ng/ml dox for 24 hours resulted in activation of *Gli1* transcript and GLI1 protein expression (GLI1 overexpression (GLI1OE)) (Figure 3B and Supplemental Figure 5A), as well as activation of the SHF Hh target gene *Foxf1* (Supplemental Figure 5B) (Hoffmann et al., 2014). Dox washout at D6 resulted in the rapid depletion of *Gli1* expression (Supplemental Figure 5F). We observed that GLI1 OE from D5-D6 caused a significant reduction in CM differentiation at D8, including a decreased percentage of cells expressing cardiac troponin (cTnT) (*p* = 0.18) (Supplemental Figure 5C), decreased relative area of cTnT-positivity (*p* = 0.03) (Figure 3D-E), and a decreased number of beating CM foci at D8 (*p* = 0.01) (Figure 3F) compared to untreated control cells. However, by D12 of the differentiation, the percentage of cTnT positive cells (*p* = 0.87), relative cTnT+expression area (*p* = 0.07), and number of beating foci (*p* = 0.13) had all become less distinguishable between GLI1 OE and control differentiations (Figure 3G-I and Supplemental Figure 5C). These observations indicated that transient activation of GLI1 in CPs delayed cardiac differentiation, but did not abrogate CP differentiation potential.

### Hh Signaling Delays Differentiation without Altering the Developmental Trajectory of Cardiac Differentiation *in vitro*

We examined the molecular basis of GLI1-induced delayed differentiation with an RNA-seq time series on GLI1 OE versus control cells. Transient activation of GLI1 (D5-6) resulted in dysregulation of 1,449 genes (log_2_ FC ≥ 0.5, FDR ≤ 0.05) at D6, including increased expression of known Hh signaling targets and SHF-expressed progenitor-specific genes *Gli1, Ptch1, Foxd1, Foxf2,* and *Osr1* (Hoffmann et al., 2014; Hynes et al., 1997; Ruiz i Altaba, 1998; Ruiz I Altaba, 1999; Sasaki et al., 1997; Vokes et al., 2008), and decreased expression of CM differentiation genes, including *Tnnt2*, *Myh6* and *Hand2* (Figure 3J and Supplemental Table 7). However, GLI1-dependent progenitor- and differentiation-specific gene expression changes normalized following dox washout at D6. The number of differentially expressed genes diminished from 1,449 at D6 to 74 by D12 (log_2_FC ≥ 0.5, FDR ≤ 0.05), consistent with the normalized CM phenotype observed at D12 (Figure 3K, Supplemental Figure 5D and Supplemental Table 7). Supporting this observation, correlation scores of GLI1 OE and control replicate transcriptomes clustered hierarchically by treatment at D6 but not at D12 (Supplemental Figure 5E). Differential expression trends over time of select CP and CM genes measured by qPCR further confirmed these findings (Supplemental Figure 5F-G).

We assessed the possibility that GLI1 OE altered the differentiation trajectories of CPs, in addition to its heterochronic effect on CM differentiation. We mapped GLI1-dependent genes onto the 19 cell type-specific clusters identified from the *in vivo* Drop-seq data to determine whether GLI1 OE promoted a non-cardiac, alternative cell fate. We observed that the top 100 GLI1 OE repressed genes were predominately expressed in the CM Drop-seq cluster (Supplemental Figure 5H), whereas the top 100 GLI1 OE activated genes were expressed in the two pSHF Drop-seq clusters (Supplemental Figure 5I), corresponding with *Gli1* expression (Supplemental Figure 5J). Furthermore, we observed that GLI1 induction *in vitro* caused gene expression changes in CPs that globally mirrored differentiation stage-specific changes *in vivo*. GLI1 OE caused global activation of pSHF-specific genes and repression of HT-specific genes, as assessed by bulk RNA-seq performed on wild type E10.5 embryos (activated vs all expressed genes *p-*value = 7.16e^−20^, repressed vs all expressed genes *p*-value = 1.49e^−57^) (Supplemental Figure 5K). These results suggested that GLI1 OE did not promote an alternative non-cardiac cell fate *in vitro*, but instead only delayed CM differentiation and prolonged their pSHF-like CP status.

We next mapped GLI1-dependent genes from mESC-CPs *in vitro* onto the Drop-seq pseudotime trajectory of gene expression *in vivo* to examine the relationship between GLI1-dependent gene expression *in vitro* and CP, intermediate, or CM stage-specific gene expression from the *in vivo* cardiac differentiation trajectory. GLI1-repressed genes sorted into two distinct clusters enriched with CM-expressed (cluster A) and intermediate-expressed (cluster B) genes, with corresponding GO terms related to heart development, heart contraction and heart conduction (Figure 3L), indicating that GLI1 OE *in vitro* repressed genes corresponding to both the drivers (*Gata4, Tbx3*) and functional products (*Ryr2, Kcnj5*) of cardiac differentiation *in vivo*. GLI1-actviated genes mapped onto the SHF Drop-seq pseudotime trajectory were sorted into four distinct clusters, with the majority expressed in the two CP-enriched clusters, including known direct Hh target genes (Figure 3M, clusters A and B). Consistently, GO terms associated with genes in these clusters identified Hh signaling itself, as well as terms associated with Hh-dependent processes such as ureteric bud development, heart development, and basal cell carcinoma. Clusters A and B were enriched for genes encoding TFs and signaling pathway components (Figure 3N), with the FOX TF family emerging as the most enriched TF family among GLI1-induced genes (Figure 3O). Together, these results confirmed that transient GLI1 OE in CPs *in vitro* maintained a CP-like expression profile reminiscent of cells in the pSHF. Furthermore, GLI1 OE *in vitro* activated a transcriptional regulatory network corresponding to a progenitor-specific, Hh-mediated network *in vivo.* We nominate this network as a heterochronic pathway preventing the onset of the CM differentiation gene expression program.

### Sustained Hh Signaling Delays Differentiation of mESC-derived Neural Progenitors

Given the widespread deployment of Hh signaling during development, we hypothesized that a heterochronic role for Hh signaling during cardiac differentiation may reflect a general differentiation timing control paradigm utilized in other developmental contexts. A Hh signaling gradient specifies distinct classes of neurons in the vertebrate neural tube and is essential for central nervous system development (Belgacem et al., 2016; Briscoe et al., 2000; Dessaud et al., 2008; Ericson et al., 1996). We investigated whether active Hh signaling also controlled neuronal differentiation timing by examining the effect of GLI1 OE during neuronal differentiation from mESCs (Supplemental Figure 6A). mESCs were directed into the neuronal lineage via biologically relevant intermediates, from neuro-ectodermal progenitors (D2-4), to neural progenitors/neurospheres (D4-6), to immature neurons with an axonal network (D6-8) (Ferreira and Caceres, 1992; Kutejova et al., 2016; Sasai et al., 2014) (Supplemental Figure 6B). Endogenous Hh signaling, reflected by *Gli1, Ptch1* and *Hhip* expression, while active throughout the course of neural differentiation, was reduced during the neural progenitor stage (Supplemental Figure 6C), suggesting that reduction of Hh signaling at this time point may be permissive for the transition of neuronal progenitors to immature neurons.

We observed that transient GLI1 OE in neuronal progenitors delayed the onset of hallmarks of neuronal differentiation *in vitro*. Dox-induced GLI1 OE from D3 to D5 caused a 25-fold increase in *Gli1* expression relative to untreated cells at D5, which subsided to baseline levels by D6 (Figure 4A-B) (Supplemental Figure 7A). In control cells, axons began to span two or more neurospheres by D5 and were marked by expression of the pan-neuronal microtubule marker tubulin β-3 (*Tubb3/*TUJ1) (Figure 4C-E) (Ferreira and Caceres, 1992), In contrast, GLI1 OE from D3-5 caused a significant reduction in axon emergence and TUJ1 immunostaining at D5 (Figure 4C-E), although the overall number of neurospheres remained unaffected (Figure 4D). By D10, however, an axon network had formed in both GLIOE and control cells and TUJ1-expression in GLI1 OE cells was statistically indistinguishable from control cells (Figure 4F-G). These observations suggested that transient GLI1 OE may delay the onset of pan-neuronal hallmarks of differentiation without altering neuronal differentiation potential, similar to its effect in CPs.

**Figure 4.**
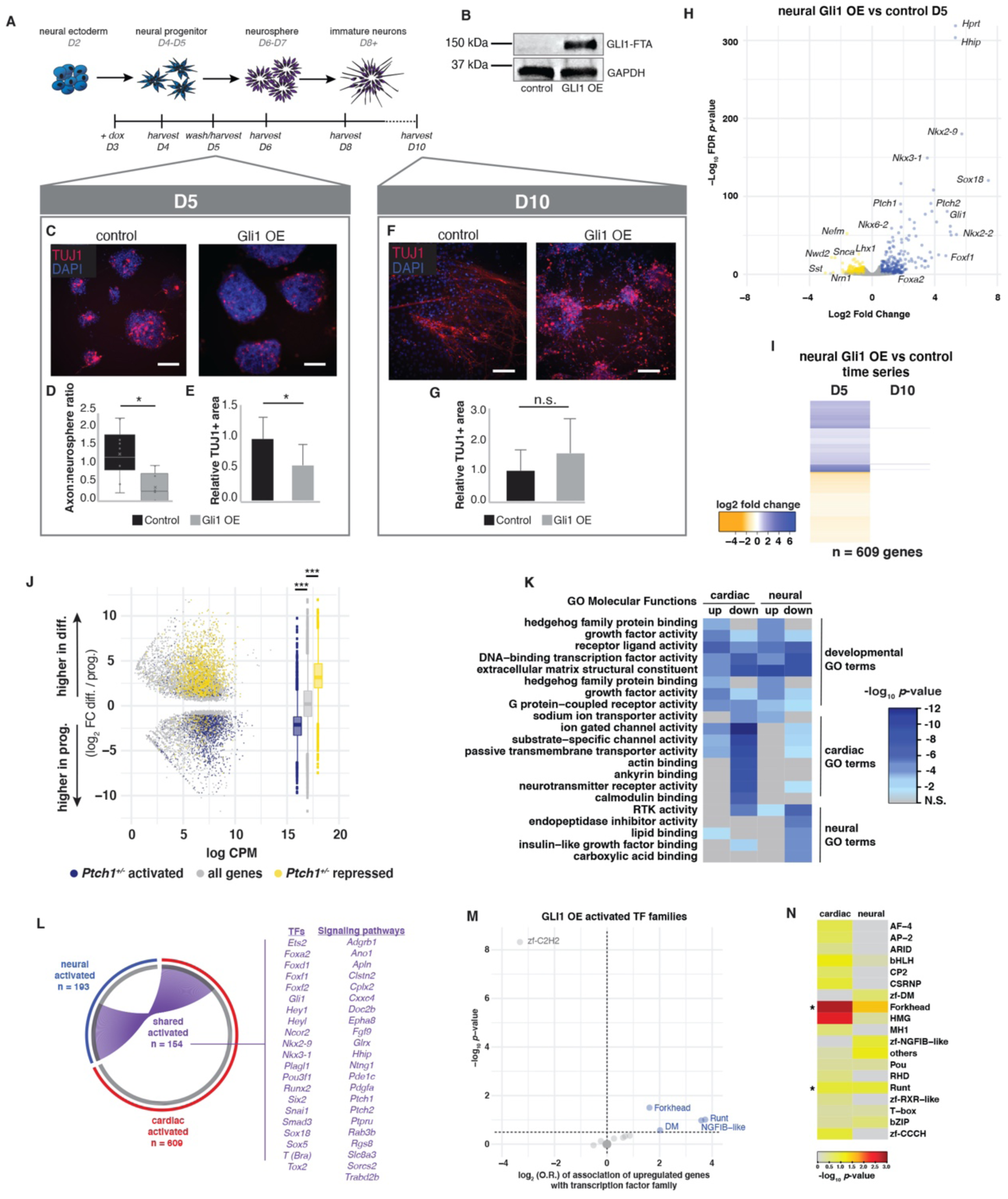
GLI^A^ expression in neural progenitors delays neuron differentiation. **(A)** Schematic representation of the transient GLI1 OE experimental design employed in mESC-derived differentiating neurons. Cells were treated with doxycycline in the neural progenitor state from D3-5, doxycycline was removed at D5, and cells were permitted to continue differentiation until the immature neuron state at D10. **(B)** Western blot showing induction of GLI1 protein in neural progenitors express after doxycycline treatment for 48 hours. **(C)** Immunofluorescent staining for pan-neuronal marker TUJ1 in control and GLI1 OE cells harvested at D5. TUJ1 staining is shown in red and DAPI counterstain is shown in blue. Scale bar = 100μm. **(D)** Quantification of the ratio of axons to neurospheres in control and GLI1 OE cells at D5. (Students T-test, * *p*-value ≦ 0.05). **(E)** Quantification of the area of TUJ1-positivity in control and GLI1 OE cells at D5. (Students T-test, * *p*-value ≦ 0.05). **(F)** Immunofluorescent staining for TUJ1 in control and GLI1 OE cells harvested at D10. TUJ1 staining is shown in red and DAPI counterstain is shown in blue. Scale bar = 100μm. **(G)** Quantification of the area of TUJ1-positivity in control and GLI1 OE cells at D10. (Students T-test, n.s. = not significant). **(H)** Volcano plot displaying activated and repressed genes in the GLI1 OE neural progenitors relative to control neural progenitors at D5. Blue and yellow dots signify significantly dysregulated genes (log_2_ FC > 0.5, FDR < 0.05). **(I)** Heatmap time-series of the log_2_ FC values of genes differentially expressed in GLI1 OE neural progenitors relative to control neural progenitors at D5 and at D10. **(J)** MA plot and box plots illustrating the distribution of *Ptch1^+/-^* dysregulated genes in the context of differential expression between ectodermal progenitors and differentiated ectoderm stages. Three *Ptch1^+/-^* activated genes that are markers of differentiated neurons are highlighted. ANOVA for three groups: *F-value* = 1557.0, *p-value* = < 2e^−16^; post-hoc Tukey Test: activated vs all genes *p-value* < 2e^−16^, repressed vs all genes p-value < 2e^−16^. **(K)** Heatmap displaying enrichment scores (log_10_ *p*-value) for GO terms associated with genes dysregulated due to GLI1 OE in cardiac and neural progenitors. Up, activated genes; down, repressed genes. **(L)** Circos plot representing the overlap of genes activated by GLI1 OE in cardiac and neural progenitors. Select shared activated genes encoding TFs or signaling pathway components are highlighted. Blue, all neural activated genes; red, all cardiac activated genes; purple, all shared activated genes. **(M)** Dot plot showing enrichment (Log_2_ odds ratio (O.R.) vs −log_10_ *p*-value) of genes encoding TFs in the Forkhead (FOX), Runt and NGFIB-like families within genes activated by GLI1 OE. **(N)** Heatmap displaying enrichment scores (log_10_ *p*-value) for TF family membership in GLI1 activated genes in cardiac and neural progenitors. Asterisks denote TF families demonstrating enrichment in both cardiac and neural progenitors (*p*-value < 0.05).

GLI1 OE in neuronal progenitors caused a temporary maintenance of neuronal progenitor gene expression at the expense of neuronal differentiation gene expression. Transcriptional profiling revealed that GLI1 OE (D3-5) caused altered expression of 523 genes in neuronal progenitors compared to control progenitors at D5, including activation of known GLI^A^ targets in the neural tube, such as *FoxA2* and *Nkx2-2* (Hynes et al., 1997, 2000; Kutejova et al., 2016; Peterson et al., 2012; Sasaki et al., 1997; Vokes et al., 2007), and repression of neuronal differentiation markers, including *Tubb3*, *Nefm*, *Sst,* and *Map2* (log_2_ FC ≥ 0.5, FDR ≤ 0.05) (Figure 4H and Supplemental Table 8). However, neural progenitor- and neuron-specific gene expression had almost entirely normalized relative to control cells by D10, as only 13 genes remained significantly dysregulated, consistent with the observed phenotypic recovery (Figure 4I, Supplemental Figure 7B-D and Supplemental Table 8). Time-course analysis of Hh-dependent gene expression revealed the transient nature of GLI1-induced differentiation delay in both cardiac and neural progenitors, demonstrating that GLI^A^ activity can function across lineages to maintain progenitor-specific gene expression and repress differentiation gene expression to effect a delay in differentiation timing.

### Maintenance of Neural Progenitor Gene Expression in Hh-dependent Medulloblastoma

As an extension of a progenitor maintenance role for Hh signaling, we hypothesized that Hh-associated neuronal cancers may be caused by the preservation of neuronal progenitor-specific gene expression with impaired differentiation. Inappropriate Hh signaling activation is a cause of medulloblastoma, a neuronal cancer that can be modelled in the adult mouse by *Ptch1* heterozygosity (*Ptch1^+/-^*) (Goodrich, 1997). *Ptch1^+/-^* mice demonstrate reduced expression of PTCH1, a critical negative feedback inhibitor of Hh signaling, resulting in abnormally high Hh pathway activity (Hahn et al., 1996; Johnson et al., 1996). We examined whether *Ptch1*-dependent gene expression in the *Ptch1^+/-^* adult mouse medulla (*Ptch1^+/-^* vs. *Ptch1^+/+^* fold change) (Grausam et al., 2017) correlated with stage-specific gene expression in the differentiating ectoderm. We identified stage-specific gene expression programs by performing differential expression analysis between ectodermal progenitors and differentiated neuronal tissues from available datasets (log_2_ FC ≥ 0.5, FDR ≤ 0.05) (Supplemental Table 9) (ENCODE Project Consortium, 2012). We observed a strong, negative correlation between expression changes due to *Ptch1* heterozygosity and expression changes during ectodermal differentiation (Supplemental Figure 7E). We compared *Ptch1*-dependent gene expression with genes differentially expressed between ectodermal progenitors and differentiated neurons. *Ptch1^+/-^-*activated genes were highly over-represented within ectodermal progenitor-specific genes and *Ptch1^+/-^-*repressed genes were over-represented within differentiated neuron-specific genes (Figure 4J), consistent with previous work examining expression changes in the postnatal mouse brain (Lee et al., 2003). Overall, our observations indicated that Hh signaling promoted progenitor-specific gene expression programs while preventing differentiation-specific gene expression in cells of diverse developmental origins and in a *Hh*-dependent cancer model.

### FOX TFs Are Enriched in Hh-dependent Cardiac and Neural Progenitor Networks

Given the similar effects of GLI1 OE in cardiac and neural progenitors, we reasoned that the Hh-induced progenitor-specific gene network might share a common effector transcription factor kernel across germ layers. Comparative Molecular Function (MF) GO term analysis on GLI1-dependent genes activated in cardiac and neural progenitors demonstrated high overlap for terms associated with activated genes and included DNA-binding TFs and signaling pathway components (Figure 4K). In contrast, MF GO terms associated with genes repressed were lineage-specific, associated with cardiac differentiation terms such as ion gated channel and actin binding in CPs and associated with neuronal functional terms such as RTK activity and endopeptidase inhibitor activity in neural progenitors, respectively. We directly compared GLI1-induced genes in cardiac and neural progenitors for overlap. Remarkably, nearly half of GLI1-induced genes in neural progenitors overlapped with GLI1-induced genes in CPs (Figure 4L), demonstrating that some specific GLI1 targets are activated across lineages. TF family enrichment analysis within GLI1-activated progenitor genes in both CPs and neural progenitors highlighted the FOX TF family as the most enriched (Figure 3O and Figure 4M). In fact, side-by-side analysis demonstrated that the FOX TF family was the only TF family highly enriched in both GLI1-activated cardiac and neural progenitor networks (Figure 4N). These observations suggested that FOX TFs may modulate the heterochronic Hh-response in diverse progenitor populations.

Hh signaling has been shown to promote proliferation in several developmental, regenerative and cancer contexts (reviewed in Jiang and Hui, 2008; Lum and Beachy, 2004; Ruiz i Altaba et al., 2007), and GLI TFs bind to regulatory elements near cell cycle genes including *Ccnd1*, *Ccnd2* and *Cdk6* genes in some contexts (Hasenpusch-Theil et al., 2018; Hu et al., 2006; Singh et al., 2018), suggesting a direct link between GLI TFs and cell cycle machinery gene expression. We reasoned that GLI1 may activate cell cycle gene expression in specified cardiac and neural progenitors as a mechanism of differentiation delay. We filtered the GO term enrichment results for GLI1-activated genes in cardiac (D6) and neural progenitors (D5) for terms associated with proliferation, cell division, the cell cycle, or mitosis, and examined the genes associated with each GO term. Proliferation-associated GO term enrichment was weak (-log_10_ *p*-value < 5) for activated genes in neural progenitors and CPs, relative to morphogenesis, differentiation and TF activity terms (Figure 3M and Figure 4K). Furthermore, we identified only one dysregulated cell cycle gene (*Ccnd1*) associated with these GO terms in neural progenitors (Supplemental Table 10) and zero in CPs (Supplemental Table 11). Instead, the genes associated with GLI1-induced proliferation GO terms in both cellular contexts were dominated by TFs (*Osr1, Foxf1, Sox11, Nr2f2*), intercellular signaling components (*Pgf, Efnb1, Fgf9, Bmp7*), and Hh pathway genes (*Gli1, Ptch1*). Together, these results failed to provide compelling evidence that GLI1 OE induced genes encoding cell cycle regulators in specified cardiac and neural progenitors, with the exception of *Ccnd1,* as a mechanism underlying differentiation delay. Instead, these results are consistent with a paradigm by which GLI1 activity in progenitors primarily induces the expression of a suite of progenitor-specific transcriptional regulators, specifically the FOX TF family, as a candidate common effector of Hh-dependent differentiation delay.

### FOXF1-mediated Repression of the Cardiac Differentiation Program

We hypothesized that the GLI1-dependent FOX TFs were functional components of the Hh-dependent heterochronic regulatory network in CPs and we therefore investigated their potential to abrogate cardiac differentiation. GLI1 OE in CPs activated the expression of eight FOX TF genes. Seven of these eight (*Foxo1, Foxd2os, Foxf1, Foxc2, Foxl1, Foxf2* and *Foxd1*) were expressed in a highly CP-specific pattern in the Drop-seq cardiac differentiation trajectory (Supplemental Figure 8A). *Foxf1* encoded the FOX TF that was expressed most specifically in the CP state and that showed the steepest decline during cardiac differentiation (Figure 5A). *Foxf1* is expressed in a Hh-dependent manner in the SHF, where it is required for cardiac morphogenesis (Hoffmann et al., 2014). We hypothesized that FOXF1 may inhibit cardiac differentiation to effect Hh-dependent differentiation delay.

**Figure 5.**
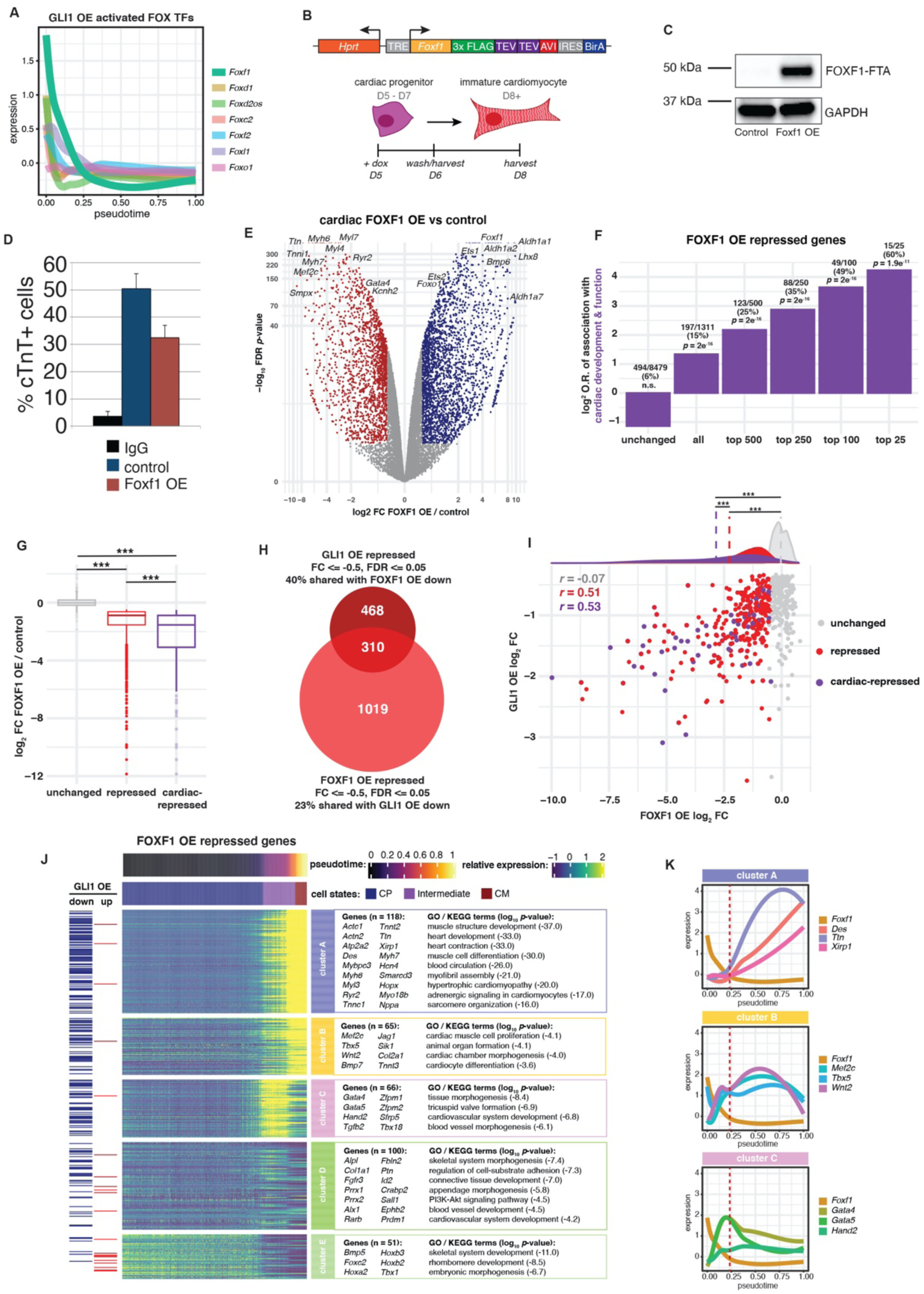
FOXF1 represses the cardiac differentiation program in CPs. **(A)** Line plot from cells in the cardiac-associated Drop-seq clusters showing the wild type expression of GLI1 OE activated FOX TFs along the pseudotime differentiation trajectory. **(B)** Diagram of the doxycycline-inducible FOXF1-FTA transgenic cassette inserted into the *Hprt* locus in mESCs, and a schematic representation of the FOXF1 OE experimental design employed in mESC-derived differentiating CMs. Cells were treated with doxycycline in the CP state from D5-6, doxycycline was removed at D6 and cells were permitted to continue differentiation until the CM state at D8. **(C)** Western blot showing induction of FOXF1 protein after doxycycline treatment for 24 hours. **(D)** Percentage of cTnT+ control and FOXF1 OE samples at D8 of differentiation, as measured by fluorescence-activated cell sorting. **(E)** Volcano plot displaying activated and repressed genes in FOXF1 OE CPs relative to control CPs at D6. Blue and red dots signify significantly dysregulated genes (log_2_ FC > 0.5, FDR < 0.05). **(F)** Bar chart showing the enrichment of cardiac-associated genes within sub-groups of all FOXF1 OE repressed genes (one-sided Fisher’s Exact Test). **(G)** Boxplot showing the log_2_ FC of unchanged genes, all repressed genes, and cardiac-repressed genes due to FOXF1 OE (Kruskal-Wallis rank sum test, χ_2_ = 3830.5, df = 2, *p*-value = 2e^−16^, followed by pairwise T-test with Benjamini-Hochberg correction, *** *p*-value ≦ 1e^−9^). **(H)** Venn diagram of intersection of GLI1 OE repressed genes and FOXF1 OE repressed genes. **(I)** Dot plot and density plot comparing the log_2_ FC of unchanged, all repressed and cardiac-repressed genes due to GLI1 OE and FOXF1 OE. Density means are shown with dotted lines. (ANOVA followed by post-hoc Tukey Test, *** *p*-value ≦ 2e^−5^). **(J)** Heatmap showing denoised data from individual cells from cardiac-associated Drop-seq clusters indicating the wild type expression levels of FOXF1 OE repressed genes along the pseudotime differentiation trajectory. Cells are ordered on the x-axis by pseudotime and are clustered on the y-axis using k-means clustering (k=5). GO terms associated with repressed genes in each cluster are also shown. Presence of gene in GLI1 OE repressed (down) or activated (up) gene sets shown in marginal rug plot. CP, cardiac progenitor; CM, cardiomyocyte. **(K)** Line plot from cells in the cardiac-associated Drop-seq clusters showing the wild type expression of *Foxf1* and select genes from clusters A, B and C of FOXF1 OE repressed genes along the pseudotime differentiation trajectory.

To examine the effect of transient FOXF1 expression on cardiac differentiation *in vitro*, we generated a dox-inducible FOXF1 OE mESC cell line (FOXF1-FTA). This line demonstrated robust FOXF1 expression at D6 from dox treatment at D5 (Figure 5B-C). We examined whether FOXF1 OE in CPs at D5 affected cardiac differentiation at D8. Similar to GLI1 OE, FOXF1 OE for 24 hours from D5-D6 caused a reduction in cTnT-positivity at D8 relative to untreated controls (∼30% vs. ∼50%) (Figure 5D). We next performed transcriptional profiling of FOXF1 OE CPs at D6 to define FOXF1-dependent gene expression in CPs. FOXF1 OE resulted in the dysregulation of 3,143 genes (FC > 0.5, FDR < 0.05) (Supplemental Table 12) including significant downregulation of many known CM differentiation products, including *Myh7* and *Ryr2* (Figure 5E and Supplemental Figure 8B-C), consistent with the ability of FOXF1 to function as a transcriptional repressor *in vivo* (Mahlapuu et al., 2001). 50% of the top 100 repressed genes were associated with cardiac development or function (hereafter referred to as cardiac-repressed genes) (Figure 5F, see Methods), and the genes most highly repressed by FOXF1 expression were cardiac-related genes (p < 2e^−16^) (Figure 5). Remarkably, FOXF1 OE repressed genes accounted for 40% of the GLI1-repressed genes (310/778) (Figure 5H), the fold change of FOXF1 / GLI1-shared cardiac repressed genes was highly correlated between GLI1 OE and FOXF1 OE treatments, and the shared FOXF1 / GLI1 co-repressed genes were highly enriched for CM differentiation and disease genes (Supplemental Figure 8D), and (Figure 5I). Together these observations indicated that FOXF1 is a critical downstream effector of cardiac differentiation delay by Hh signaling in CPs.

We investigated the dynamic relationship between FOXF1-dependent gene expression *in vitro* and cardiac differentiation *in vivo*, mapping FOXF1 OE-repressed genes onto the Drop-seq cardiac differentiation trajectory. FOXF1 OE-repressed genes were sorted into 5 major pseudotime clusters (Figure 5J). Most FOXF1 OE repressed genes that were expressed at some point along the pseudotime differentiation trajectory (249/401, 62%) were either highly expressed only during late cardiac differentiation (Cluster A), highly expressed during late cardiac differentiation with activation in the intermediate stage (B), or highly expressed during the intermediate stage (Cluster C), a property shared with the majority of GLI1 OE repressed genes (Figure 5J). Genes highly expressed in the late cardiac differentiation (cluster A) were dominated by cardiac differentiation products, including *Myh6, Tnnt2, Ttn,* and *Nppa*. Genes highly expressed during late cardiac differentiation but with activation in the intermediate stage (cluster B) included cardiogenic TFs that promote cardiac differentiation (e.g. *Mef2c* and *Tbx5*) and some cardiac differentiation products (e.g. *Tnnt3*). Several genes expressed highly in the intermediate state (cluster C) showed initial activation in the CP state, including cardiogenic TFs required for cardiac morphogenesis, such as *Gata4, Gata5* and *Hand2,* and were associated with GO terms affiliated with cardiac development as well as cardiac differentiation (Figure 5J). These results suggested that FOXF1 may inhibit cardiac differentiation by repression at multiple levels of the cardiac differentiation program. Interestingly, the genes in these clusters demonstrated distinct expression dynamics relative to FOXF1 expression. Genes in cluster A were only activated following the near-complete depletion of FOXF1, while genes in clusters B and C demonstrated activation prior to depletion of FOXF1 expression (Figure 5K). These observations indicated that repressed genes exhibited variable sensitivities to FOXF1 levels. Genes in clusters D and E showed weaker expression across pseudotime and were associated with developmental processes other than cardiac differentiation, suggesting that FOXF1-mediated repression may be deployed in diverse lineages (Figure 5J). Overall, these results implicated FOXF1 in the repression of the pro-cardiac differentiation network.

### FOXF1 Directly Represses a Cardiomyocyte Differentiation Program

We hypothesized that FOXF1 repressed cardiac differentiation by directly inhibiting cardiac differentiation gene expression. We identified FOXF1 binding sites genome-wide by ChIP-seq for FOXF1 in GLI1 OE CPs (Figure 6A). We identified 8,398 consensus FOXF1 binding sites, including at the known FOXF1 target gene *Pecam1* (Ren et al., 2014) and near its own promoter (Figure 6B, Supplemental Figure 9A-B and Supplemental Table 13). A FOXF1 DNA-binding motif was the most highly enriched TF motif and was present at 77% of FOXF1-localized regions (Supplemental Figure 9C). To identify genes directly regulated by FOXF1, we analyzed the location of FOXF1 binding sites with respect to genes dysregulated by FOXF1 OE. We observed that FOXF1 binding was enriched near the TSSs of cardiac-repressed genes compared to all repressed genes or unchanged genes (Figure 6C). 90% of cardiac-repressed genes had a FOXF1 binding site within 250kb of the TSS, and cardiac-repressed genes with FOXF1 binding within 250kb exhibited significantly stronger repression compared to all repressed genes (Supplemental Figure 9D). FOXF1 OE cardiac-repressed genes with FOXF1 binding within 250kb were enriched for cardiac GO molecular function (MF) terms including contraction (GO:0003779, actin binding) and electrophysiology (GO:0046873, metal ion transporter activity) as well as transcriptional regulation (GO:0008134, transcription factor binding) (Supplemental Figure 9E). The most dramatic repression was observed for genes associated with CM contraction (Figure 6D), such as *Mybpc1*, the most highly repressed FOXF1-bound cardiac-repressed gene, and *Tnnt2*. These results suggested that FOXF1 directly targets and represses genes essential for cardiomyocyte differentiation.

**Figure 6.**
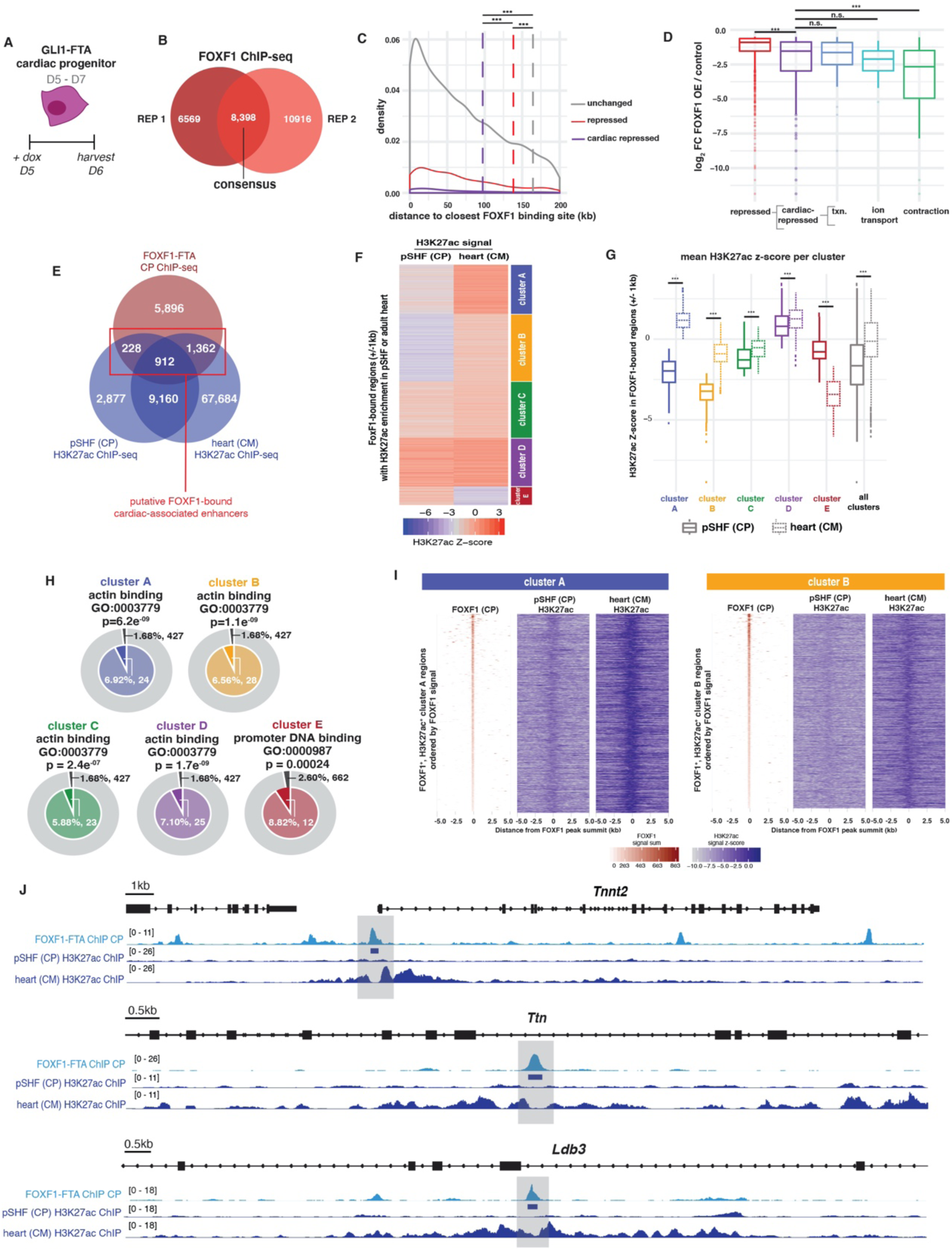
FOXF1 directly represses the cardiac differentiation program by binding to cardiomyocyte gene enhancers. **(A)** A schematic representation of FOXF1 ChIP-seq experimental design in CPs derived from the GLI1-FTA mESC line. Cells were treated with doxycycline in the CP state from D5-6 and harvested at D6. **(B)** Venn diagram showing the number of consensus FOXF1 binding regions found between two biological replicates. **(C)** Density plot displaying the mean distance of FOXF1 OE unchanged, all repressed and cardiac-repressed gene sets from the nearest FOXF1 binding site (Kruskal-Wallis rank sum test, χ_2_ = 37.243, df = 2, *p*-value = 8.179e^−9^, followed by pairwise T-test with Benjamini-Hochberg correction, *** *p*-value ≦ 0.001). **(D)** Boxplots of the FOXF1 OE / control log_2_ FC of sub-groups of FOXF1 cardiac-repressed genes based on the associated molecular function of the encoded protein (Kruskal-Wallis rank sum test, χ_2_ = 103.86, df = 4, *p*-value = 2e^−16^, followed by pairwise T-test with Benjamini-Hochberg correction, *** *p*-value ≦ 0.001). **(E)** Venn diagram of FOXF1 binding sites intersected with regions enriched for H3K27ac histone modification in CPs from the pSHF and CMs from the adult heart. H3K27ac+ regions that overlap with FOXF1 binding sites were considered putative FOXF1-bound cardiac enhancers. **(F)** Heatmap showing the H3K27ac signal Z-scores in CPs and CMs for FOXF1-bound regions. Regions were clustered using k-means clustering (k=5). **(G)** Boxplots comparing the H3K27ac Z-scores of FOXF1-bound regions in clusters A-E between pSHF/CP and adult heart/CM states (ANOVA, followed by Mann-Whitney-Wilcoxon post-hoc test, \*\*\**p*-value ≦ 2.1e^−12^). **(H)** Nested pie-charts demonstrating enrichment of genes associated with actin binding (clusters A-D) or DNA binding (cluster E) within FOXF1 OE repressed genes associated with FOXF1-bound regions in clusters A-E. **(I)** Heatmaps showing FOXF1 binding and H3K27ac signal z-scores in CPs and CMs for FOXF1-bound regions in clusters A & B. **(J)** Genome browser views of three FOXF1-bound regions within genes encoding proteins required for CM contraction. FOXF1 and CP / CM H3K27ac fold-enrichment ChIP signal over input is shown, and FOXF1-bound regions are highlighted in grey.

We evaluated the enhancer dynamics of FOXF1 binding sites to investigate the mechanism by which FOXF1 binding in CPs inhibits the expression of cardiac differentiation genes. We first identified and compared the cardiac enhancer landscape of CPs and CMs *in vivo* by analyzing the localization of H3K27ac, a histone modification observed at active enhancers (Creyghton et al., 2010). We performed ChIP-seq for H3K27ac in CPs in the pSHF of E9.5 embryos, identifying 13,177 locations of H3K27ac enrichment, including at the promoters of CP genes, like *Isl1,* and not at the promoters of cardiomyocyte genes, like *Nppa* (Supplemental Figure 9D). ChIP-seq for H3K27ac+ in the adult heart previously identified 79,118 locations of enrichment (Yue et al., 2014). From the CP/CM union set of 82,223 H3K27ac enriched cardiac enhancers, 3,105 were CP-specific, 69,046 were CM-specific, and 10,072 were shared. We analyzed FOXF1 binding sites (summit +/− 1kb) with H3K27ac enriched regions, and observed that 30% (2,508/8,398) of FOXF1 binding sites were associated with cardiac enhancers (Figure 6E). H3K27ac enrichment at FOXF1-bound cardiac enhancers was generally stronger in the CM than the CP state (Supplemental Figure 9G). Clustering of FOXF1-bound cardiac enhancers revealed five enhancer clusters (Clusters A-E), of which four (Clusters A-D) demonstrated significantly increased in H3K27ac enrichment in the CM relative to the CP state (Figure 6F-G). These four clusters harbored the vast majority of FOXF1 bound enhancers (92%; 2,316/2,508). The remaining cluster (Cluster E) demonstrated higher H3K27ac enrichment in the CP state (Figure 6F-G). The dynamic pattern of H3K27ac defined predominantly cardiac differentiation enhancers bound by FOXF1 in CPs that acquired activity during the transition from CP to CM, following FOXF1 depletion. These observations suggested that FOXF1 directly repressed cardiac differentiation gene expression.

We assessed the regulatory effect of FOXF1 binding at cardiac enhancers by linking the closest TSS of FOXF1 OE repressed genes to the clustered FOXF1-bound cardiac enhancers. The most highly enriched GO molecular function term for genes linked to FOXF1-bound cardiac enhancers that acquired H3K27ac signal in CMs (clusters A-D) was actin binding (Figure 6H). Two clusters in particular, clusters A and B, demonstrated a dramatic increase in H3K27ac deposition during cardiac differentiation, and these FOXF1-bound cardiac enhancers were enriched near cardiac genes that were repressed by FOXF1 OE (Figure 6I and Supplemental Figure 9H). These FOXF1 OE cardiac-repressed genes with FOXF1 binding included *Tnnt2*, *Ttn* and *Ldb3*, three genes essential for sarcomere organization and contraction, and H3K27ac deposition at their local enhancers occurred at nucleosomes immediately adjacent to FOXF1 binding sites (Figure 6J). Binding of FOXF1 to loci that demonstrate FOXF1-dependent repression of CM gene expression provided a heterochronic mechanism by which FOXF1 binds and silences cardiac differentiation regulatory elements in CPs to delay the onset of cardiac differentiation gene expression.

### A Hh Heterochronic Network is Required to Prevent Premature Differentiation in pSHF CPs and CHD

We hypothesized that the Hh signaling heterochronic GRN we defined in CPs *in vitro* mediated the differentiation status of SHF CPs *in vivo* and was required for the morphogenesis of SHF-derived cardiac structures, including the atrioventricular septum (Briggs et al., 2016; Goddeeris et al., 2008; Hoffmann et al., 2009, 2014). To test this hypothesis, we disrupted the Hh signaling pathway at three levels of Hh signal transduction in the SHF *in vivo* - the SHH ligand, the GLI TFs and the GLI1 target effector FOXF1 - and assessed the effect on SHF differentiation timing and cardiac morphogenesis. To disrupt *Shh*, we evaluated *Shh^-/-^* (*Shh* KO) embryos. To override the wild type GLI^A^/GLI^R^ ratio in the pSHF in favor of GLI^R^, we employed Cre-dependent expression of the truncated, repressive form of GLI3 (GLI3^R^ OE) activated by a tamoxifen-inducible Cre recombinase expressed from the *Osr1* locus, a pSHF-specific gene (*Osr1^eGFPCre-ERT^*^2*/+*^; *ROSA26^Gli3R-Flag/+^*) (Mugford et al., 2008; Vokes et al., 2008). To disrupt *Foxf1*, we conditionally ablated *Foxf1* (*Foxf1* cKO) in cells receiving Hh signaling using a tamoxifen-inducible Cre recombinase expressed from the *Gli1* locus (*Gli1^Cre-ERT2/+^*; *Foxf1^fl/fl^*) (Ahn and Joyner, 2004).

Disruption of Hh signaling at all three levels of signal transduction caused precocious cardiomyocyte differentiation in the SHF. We evaluated myocardial differentiation in Hh pathway mutant embryos by immunofluorescence for sarcomeric myosin (MF20) at E10.5. Littermate control embryos demonstrated MF20 staining only in the heart, including the atrial walls, ventricles and outflow tract, but not in the SHF. In contrast, all three Hh-mutant models demonstrated ectopic MF20 expression that extended beyond the heart and into the SHF (Figure 7A, yellow arrowheads). *Shh* KO embryos demonstrated broad ectopic expression of MF20 in the SHF along the AP axis compared to control embryos, consistent with a requirement for Hh signaling in both the pSHF and aSHF (Goddeeris et al., 2008; Hoffmann et al., 2009). *Gli3^R^* OE embryos, but not littermate controls, displayed MF20 expression in the dorsal mesenchymal protrusion (DMP), a pSHF region normally occupied by undifferentiated, mesenchymal progenitor cells which migrate into the heart to form the atrial portion of the atrioventricular septum (AVS) (Figure 7A, B) (Hoffmann et al., 2009). *Foxf1* cKO, but not control embryos, demonstrated MF20 staining in the SHF that extended from the outflow tract into the dorsal mesothelium (also known as the dorsal mesocardium), a structure that contributes to the primary atrial septum (PAS) (Franco et al., 2000). These results demonstrated a requirement for Hh signaling and the Hh target gene *Foxf1* in the SHF for preventing precocious differentiation of SHF CPs.

**Figure 7.**
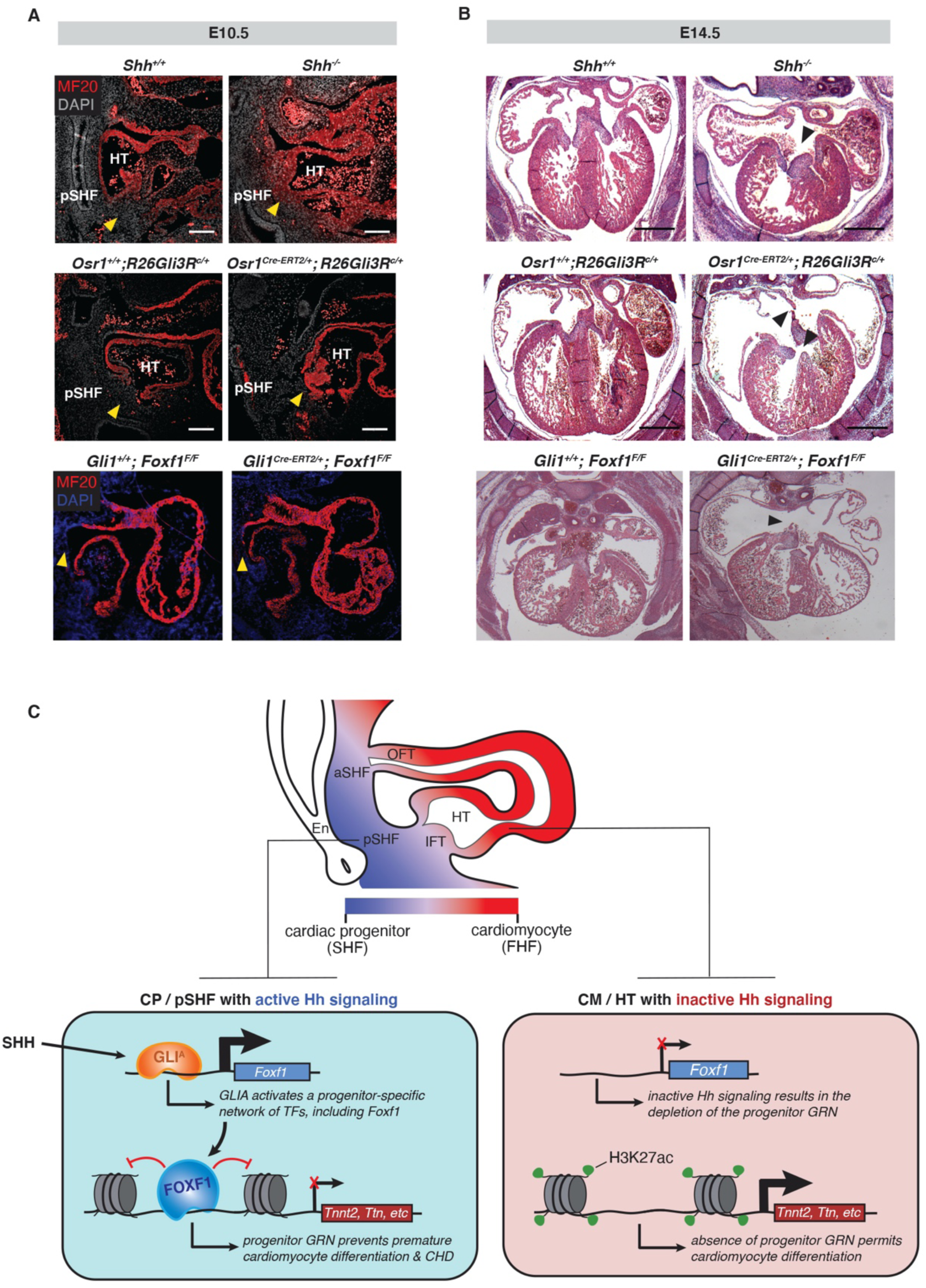
Disruption of the Hh signaling pathway at three levels causes precocious differentiation and AVSDs *in vivo*. **(A)** Immunofluorescent staining for MF20 (sarcomeric myosin) in the E10.5 pSHF of control and Hh pathway mutant embryos. MF20 staining is shown in red and DAPI counterstain is shown in grey or blue. Yellow arrowheads demarcate mesenchymal CPs migrating into the HT at the posterior aspect of the inflow tract (control) or inappropriate expression of sarcomeric myosin in the same region (Hh pathway mutants). Scale bars = 200μm. **(B)** Histological sections of E14.5 hearts from control and Hh mutant embryos. Black arrowheads indicate incidence of AVSD. Scale bars = 200μm. **(C)** Model of the Hh signaling-dependent activation of a progenitor regulatory network through GLI^A^ TFs, such as GLI1. GLI1 activates FOXF1 and other progenitor-specific TFs, which directly repress the expression of CM differentiation genes in the CP state via inhibition of the activity of cardiac enhancers. Upon removal of the Hh signal in differentiating CMs, FOXF1 expression is depleted, permitting the activation of cardiac enhancers, the expression of cardiac genes and CM differentiation.

We reasoned that premature SHF differentiation at E10.5 may disrupt AVS morphogenesis. We therefore examined each Hh pathway mutant at E14.5, when atrial septation is normally complete. *Shh^-/-^* embryos exhibited a complete loss of the atrial septum resulting in AVSDs (Figure 7B), as previously described (Izraeli et al., 1999; Kim et al., 2001; Meyers and Martin, 1999; Tsukui et al., 1999). *Gli3^R^* OE embryos exhibited AVSDs due to a diminished DMP, defects in the PAS and failure of the ventricular septum to fuse to the atrioventricular cushions. *Foxf1* cKO embryos completely lacked a PAS and DMP, also resulting in an AVSD. Taken together, these findings indicate that Hh signaling causes activation of a heterochronic network of progenitor-specific TFs, including FOXF1, that is required to actively maintain SHF CP status, and prevent premature cardiac differentiation and CHD.

## DISCUSSION

We provide evidence that Hh signaling activates a heterochronic GRN that controls differentiation timing without altering developmental potential in diverse contexts. Modeling active Hh signaling via GLI1 expression prevents differentiation of lineage-specific progenitors in cardiac and neural lineages, whereas removal of GLI1 permits differentiation of both lineages *in vitro*. We identify FOX TFs as candidate effectors of differentiation timing, and show that deployment of FOXF1 within CPs is sufficient to delay cardiac differentiation *in vitro*. We identify a cardiac differentiation GRN that is directly repressed by FOXF1 binding, as the molecular mechanism of FOXF1-mediated differentiation delay. Finally, we demonstrate that both active Hh signaling and *Foxf1* expression are required *in vivo* to prevent precocious CP differentiation and CHD. In summary, Hh signaling activates a heterochronic TF module, including FOXF1, which directly represses the cardiac differentiation GRN.

We describe a Hh-dependent two-step mechanism, in which (1) GLI1 directly activates the expression of a transcriptional repressor, which (2) directly represses a lineage-specific pro-differentiation GRN, at regulatory elements of differentiation-specific genes (Figure 7C). Our studies indicate that the pro-differentiation GRN is poised to activate cardiac differentiation gene expression if not directly bound and repressed by Hh-dependent transcriptional repressors. This transcriptional architecture provides a dominant layer of transient, precise Hh signal-dependent temporal control on top of the cardiac pro-differentiation GRN. The role of FOXF1 in the regulation of histone acetylation at cardiac differentiation genes is consistent with a transient epigenetic pausing mechanism in which removal of the Hh signal, and the Hh-dependent repressive GRN including FOXF1, enables the activation of regulatory elements of cardiomyocyte-specific genes without further intervention. An enhancer “priming” role, in which a FOX TF binds and inhibits the activation of a regulatory element required at a subsequent developmental stage, has been described for FOXD3 in pluripotent stem cells (Krishnakumar et al., 2016; Liber et al., 2010; Respuela et al., 2016). This activity may represent a generalizable function of FOX TFs with repressive activity, and is consistent with a heterochronic role for Hh-dependent FOX TFs in the delay of developmental transitions (Buecker et al., 2014; Zaret and Carroll, 2011).

We postulate that the Hh signal-induced dominant repressor GRN could be layered on diverse lineages for differentiation timing control, providing a modular system for the evolution of structurally complex organs. In the case of cardiac development, delayed differentiation of the SHF relative to the FHF, a defining distinction between the SHF and FHF, is at least in part a consequence of SHF Hh signaling (Figures 2 and 7). In Hh mutant embryos, the SHF undergoes precocious differentiation and cardiac structures normally formed by the SHF, including the dorsal mesenchymal protrusion and primary atrial septum in the posterior and pulmonary outflow tract in the anterior, fail to form (Briggs et al., 2016; Goddeeris et al., 2008; Hoffmann et al., 2009). Interestingly, CHD-causing mutations have been identified in many genes encoding components of the cilium, a cellular organelle essential for Hh signaling (Friedland-Little et al., 2011; Huangfu et al., 2003; Ocbina et al., 2011; Ozanna Burnicka-Turek, 2016; Watkins et al., 2019). These observations suggest that precocious progenitor differentiation, potentially as a result of abrogated Hh signaling, may be a significant mechanism underlying CHD.

The observation that Hh signaling can delay differentiation in both cardiac and neural lineages suggests that Hh signaling may act globally to control differentiation timing. A corollary of this hypothesis is that mutations in Hh signaling components, or cilia genes required for Hh signaling, may cause structural birth defects by allowing precocious differentiation across lineages. Hh signaling has been implicated in the development of many mammalian organs, including the palate, limbs, genitourinary tract, gastrointestinal tract, respiratory system, skeletal system, central nervous system, skin, inner ear, teeth and eye (reviewed in Briscoe and Thérond, 2013; Dworkin et al., 2016; Groves and Fekete, 2012; Haraguchi et al., 2019; Hosoya et al., 2020; Ingham and Placzek, 2006; Ingham et al., 2011; Jiang and Hui, 2008; Lee et al., 2016; McMahon et al., 2003; Singh et al., 2015; Wallace, 2008). Global disruption of Hh signaling during embryogenesis causes pleiotropic developmental syndromes, including VACTERL syndrome, Gorlin Syndrome, holoprosencephaly and Greig cephalopolysyndactyly syndrome (Hui and Angers, 2011; McMahon et al., 2003; Ngan et al., 2012; Nieuwenhuis and Hui, 2005). Elucidation of the Hh-dependent GRNs in these distinct developmental contexts will determine whether disrupted heterochronic control of progenitor differentiation timing is a molecular mechanism underlying Hh-dependent birth defects across organs.

Hh signaling has been affiliated with the progenitor cell state in developmental, adult stem cell and cancer contexts, consistent with a role as a heterochronic differentiation timing switch. Hh signaling has been implicated in multiple organ-specific progenitor functions during development, including cell migration and proliferation (Agathocleous et al., 2007; Ingham and Placzek, 2006; Jiang and Hui, 2008; Kaldis and Richardson, 2012; Liu and Ngan, 2014). In adults, Hh signaling has been implicated in the maintenance of organ-specific stem cells required for organ homeostasis (Beachy et al., 2004; Petrova and Joyner, 2014; Roy and Ingham, 2002). For example, Hh signaling promotes proliferation of neural stem cells in the adult hippocampus (Ahn and Joyner, 2005; Lai et al., 2003; Li et al., 2013; Machold et al., 2003). A role in progenitor maintenance is also consistent with the importance of Hh signaling in the regeneration of many organs, ranging from the heart (Kawagishi et al., 2018; Singh et al., 2018; Wang et al., 2015), bladder (Shin et al., 2011), prostate (Karhadkar et al., 2004), bone (Miyaji et al., 2003), tooth (Zhao et al., 2014), liver (Ochoa et al., 2010), to the lung (Peng et al., 2015; Watkins et al., 2003). Whereas controlled Hh signaling in adult stem cells has been associated with organ homeostasis and regeneration, uncontrolled Hh activation promotes cancer, including medulloblastoma, basal cell carcinoma, and rhabdomyosarcoma (Hooper and Scott, 2005; Jiang and Hui, 2008; Ng and Curran, 2011; Taipale and Beachy, 2001; Wicking et al., 1999). Progenitor-specific roles for Hh signaling have engendered the hypothesis that Hh signaling in cancer functions through promotion of cancer stem cells (Beachy et al., 2004; Ruiz i Altaba et al., 2002; Teglund and Toftgård, 2010). In most of these contexts, the transcriptional networks activated by the GLI TFs remain to be elucidated. Therefore, whether the specific control of progenitor status is reflective of direct or indirect control by TFs downstream of Hh signaling is unknown. In some contexts, direct control of the expression of cell cycle regulators by GLI TFs has been demonstrated (Hasenpusch-Theil et al., 2018; Hu et al., 2006; Singh et al., 2018). However, our results do not support a proliferation-mediated mechanism of differentiation delay in specified cardiac and neural progenitors, as GLI1-activated genes are not enriched with these gene categories. Given the similarity between our findings in CPs, neural progenitors and a medulloblastoma model, we instead nominate a heterochronic Hh-activated GRN comprised of TFs that maintain progenitor status and prevent differentiation as a plausible molecular mechanism underlying the association of Hh signaling with the progenitor cell state across developmental and disease contexts.

How the heterochronic role of Hh signaling in differentiation timing control intersects with the well-documented role of Hh signaling in tissue patterning remains an open question. A Hh-dependent GRN has been unveiled in some developmental contexts, including the limb and neural tube, where GLI^A^-dependent TFs play a direct role in tissue patterning (Lei et al., 2006; Oosterveen et al., 2012; Peterson et al., 2012; Sasaki et al., 1997; Vokes et al., 2007, 2008). Although a Hh-dependent GRN that controls patterning does not preclude a coincident, distinct Hh-dependent GRN that controls differentiation timing, a number of observations suggest that the two progenitor phenomena may be linked. As in the pSHF, GLI^A^ TFs in the neural tube and limb induce the expression of a suite of TFs with repressive activity, including FOXA2, NKX2-2, and PRDM1/BLIMP-1 (Peterson et al., 2012; Vokes et al., 2008). The repressive activity of these TFs has been implicated in cell-type specification, mechanistically by repressing transcriptional drivers of alternative cell fates. However, the repressive function of NKX2-2 in the neural tube extends to effector genes of differentiation (Balaskas et al., 2012; Kutejova et al., 2016), mirroring our observations during cardiac differentiation. Further, constitutive Hh signaling activation in the neural tube maintains these cells in a progenitor state (Rowitch et al., 1999), consistent with our observations for mouse medulloblastoma. Finally, recent transcriptomic analyses of sub-domains of the neural tube and limb show that the populations with the highest Hh activity sustain high expression of progenitor-specific genes relative to Hh-deficient domains (Delile et al., 2019; Reinhardt et al., 2019). Our results demonstrating activation of progenitor genes at the expense of differentiation genes upon transient GLI1 OE in progenitors, and the eventual phenotypic and transcriptomic recovery of cells following GLI1 removal, suggest a direct role for GLI-dependent networks in regulating differentiation timing, rather than patterning. Dissection of the morphogenic versus differentiation timing functions of Hh signaling will require further investigation of the molecular activity of Hh-dependent transcriptional regulators in each context.

The concept of heterochrony was developed to describe the evolution of structural differences between species and was defined as altered timing in the appearance of features during ontogeny (Smith, 2003). Heterochrony may play a role in the evolution of heart development. Features generated by the FHF, such as the left ventricle, are evolutionarily ancient compared to features generated by the SHF, such as the right ventricle, pulmonary outflow tract, and the atrial septum (Kelly, 2012). SHF-derived structures are predominately required for the efficient handling of pulmonary circulation. In fact, the lungs are the source of Hh ligand received by the SHF CPs and are thereby required for their development (Goddeeris et al., 2008; Hoffmann et al., 2009). Differentiation timing controlled by cell non-autonomous Hh signaling between the developing lungs and heart provides a plausible mechanism for inter-organ coordination of heart and lung development. Cardiac evolution may have employed Hh signaling as a heterochronic mechanism in lunged animals, delaying SHF differentiation to enable morphogenesis of the cardiovascular structures required for pulmonary circulation. We speculate that Hh-dependent heterochronic GRNs may have been broadly deployed to delay lineage-specified progenitor differentiation and support the evolution of complex organ development and homeostasis.

## SUPPLEMENTAL FIGURES

**Supplemental Figure 1.**
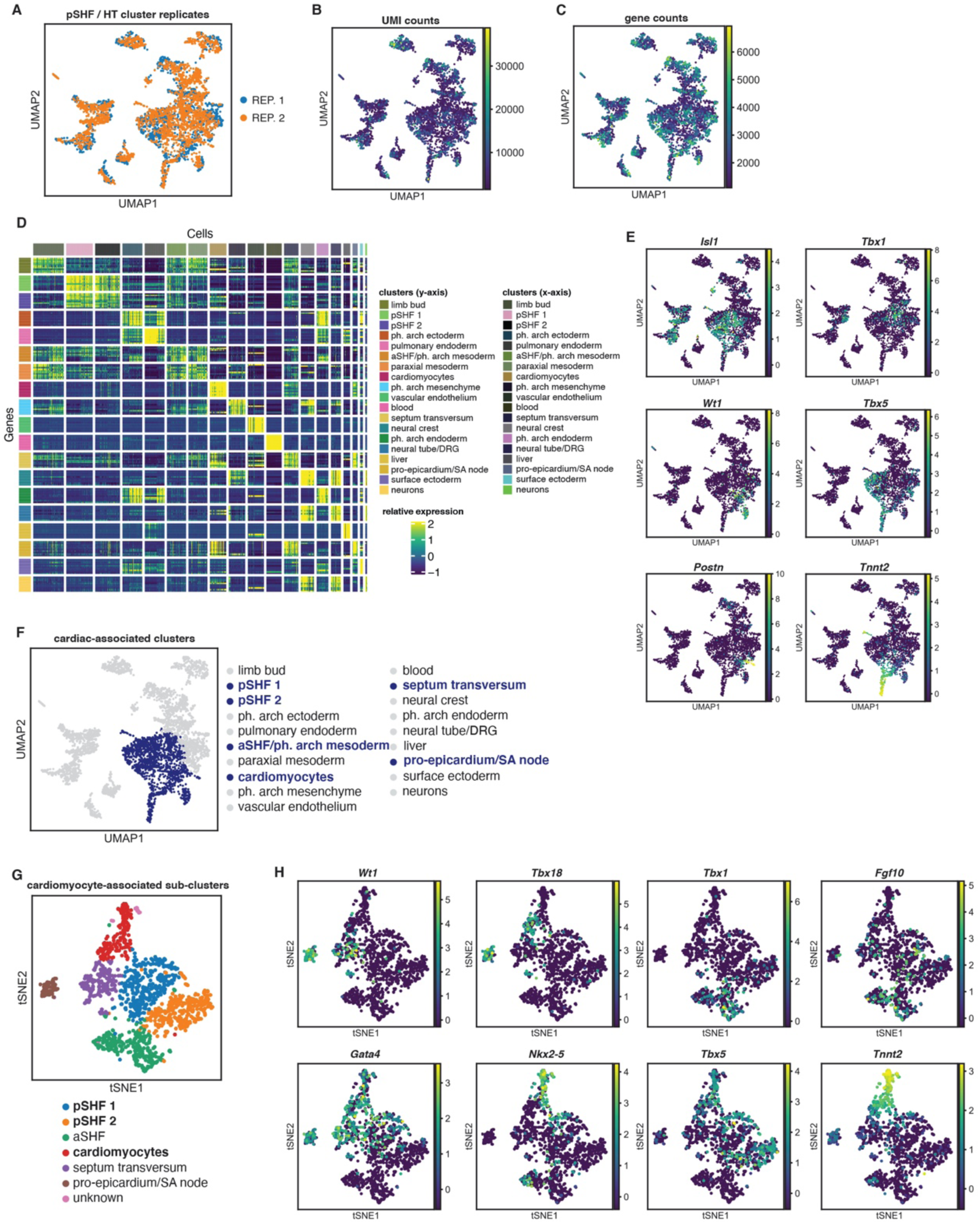
Drop-seq analysis of the wild-type E10.5 SHF and HT identifies cardiac progenitors and cardiomyocytes. **(A)** UMAP plot of biological replicate sample overlap in 19 distinct cell type clusters identified from microdissected SHF and HT tissue. **(B)** UMAP plot of UMI counts in 19 cell type clusters. **(C)** UMAP plot of gene counts in 19 cell type clusters. **(D)** Heatmap of the top 10 most-differentially expressed markers in 19 cell type clusters. Cells (x-axis) and differentially expressed genes (y-axis) are grouped by cluster ID. **(E)** UMAP plot of SHF/HT clusters indicating the expression levels of marker genes expressed in cardiac-associated clusters. **(F)** UMAP plot of SHF/HT clusters with only populations identified as cardiac-associated highlighted. **(G)** tSNE plot showing clustering of 7 CM-associated cluster populations. **(H)** tSNE plot of CM-associated clusters indicating the expression levels of marker genes expressed in CMs and various CP populations.

**Supplemental Figure 2.**
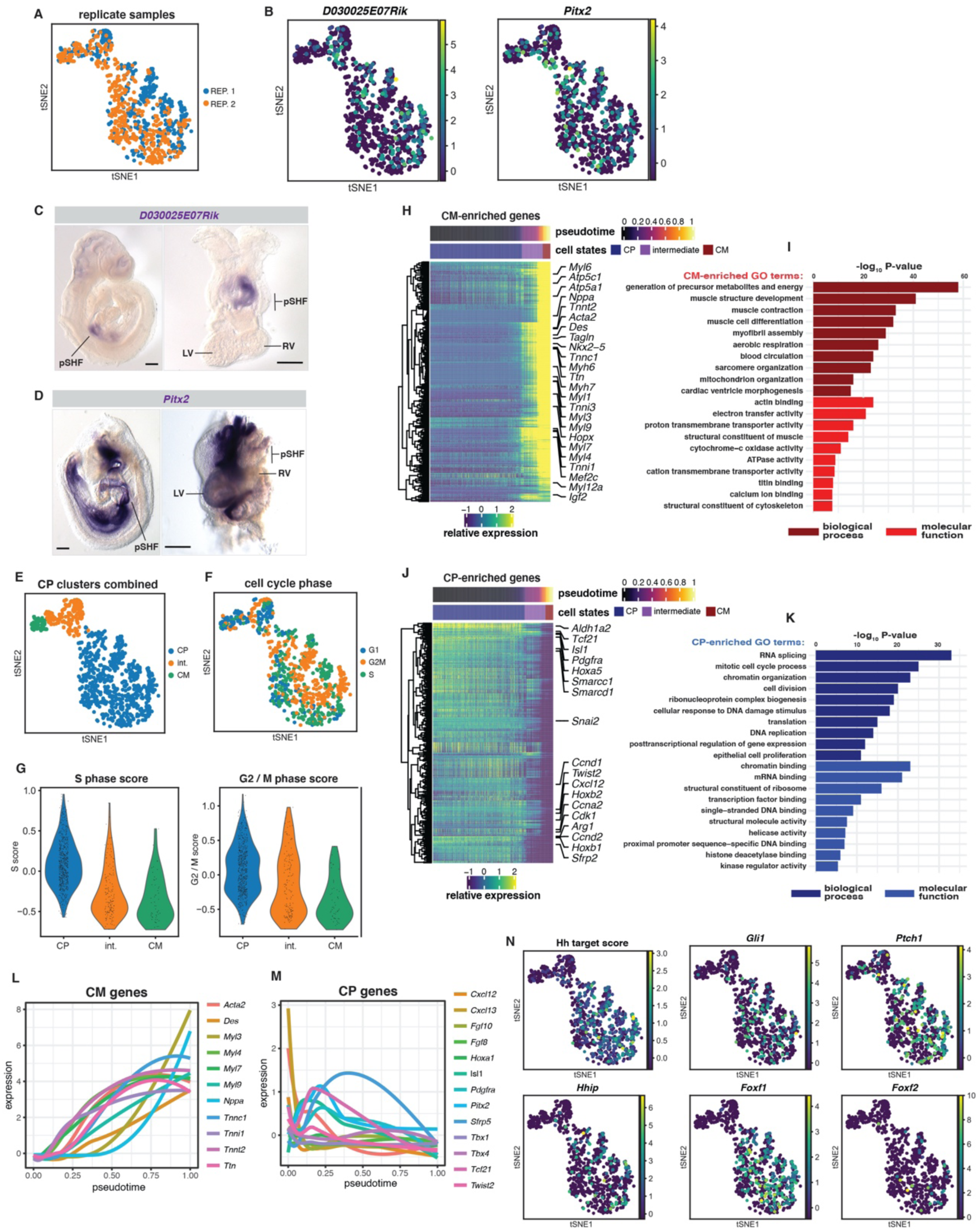
Drop-seq analysis of wild-type E10.5 pSHF and HT sub-populations reveals a cardiomyocyte differentiation continuum. **(A)** tSNE plot of biological replicate sample overlap in CM-associated clusters. **(B)** tSNE plots indicating the expression levels of side-specifc markers of SHF CPs. *D030025E07Rik/Plyrr* is expressed on the right side (CP1) and *Pitx2* are expressed on the left side (CP2). **(C)** *In situ* hybridization demonstrating the right side-specific expression domain of *D030025E07Rik/Plyrr* in the SHF of wild type E9.5 embryos. Scale bar = 100μm. The entire embryo is shown on the left, and a microdissected portion of the embryo containing the HT and pSHF is shown on the right from a posterior perspective. pSHF, posterior second heart field; LV, left ventricle; RV, right ventricle. **(D)** *In situ* hybridization demonstrating the left-side specific expression domain of *Pitx2* in the SHF of wild type E9.5 embryos. Scale bar = 100μm. The entire embryo is shown on the left, and a microdissected portion of the embryo containing the HT and pSHF is shown on the right from a posterior perspective. pSHF, posterior second heart field; LV, left ventricle; RV, right ventricle. **(E)** tSNE plot of CM-associated clusters, with left/right CP clusters combined. CP, cardiac progenitor; CM, cardiomyocyte. **(F)** tSNE plot of cell cycle phase in CM-associated clusters. **(G)** Violin plot of S phase and G2/M phase scores in CM-associated clusters. CP, cardiac progenitor; CM, cardiomyocyte. **(H)** Heatmap showing denoised data from individual cells from cardiac-associated Drop-seq clusters indicating the expression levels of genes enriched in the CM cluster, relative to the combined CP cluster, along the pseudotime differentiation trajectory. Cells are ordered on the x-axis by pseudotime and are hierarchically clustered on the y-axis. Select CM-enriched genes are highlighted. CP, cardiac progenitor; CM, cardiomyocyte. **(I)** Gene ontology (GO) analysis of CM-enriched genes. **(J)** Heatmap showing denoised data from individual cells from cardiac-associated Drop-seq clusters indicating the expression levels of genes enriched in the combined CP cluster, relative to the CM cluster, along the pseudotime differentiation trajectory. Cells are ordered on the x-axis by pseudotime and are hierarchically clustered on the y-axis. Select CP-enriched genes are highlighted. CP, cardiac progenitor; CM, cardiomyocyte. **(K)** Gene ontology (GO) analysis of CP-enriched genes. **(L)** Line plot indicating the relative expression levels of known HT/CM marker genes along the pseudotime differentiation trajectory. **(M)** Line plot indicating the relative expression levels of known pSHF/CP marker genes along the pseudotime differentiation trajectory. **(N)** tSNE plots indicating the expression levels of individual markers of active Hh signaling and a Hh target metagene score in CM-associated cell clusters.

**Supplemental Figure 3.**
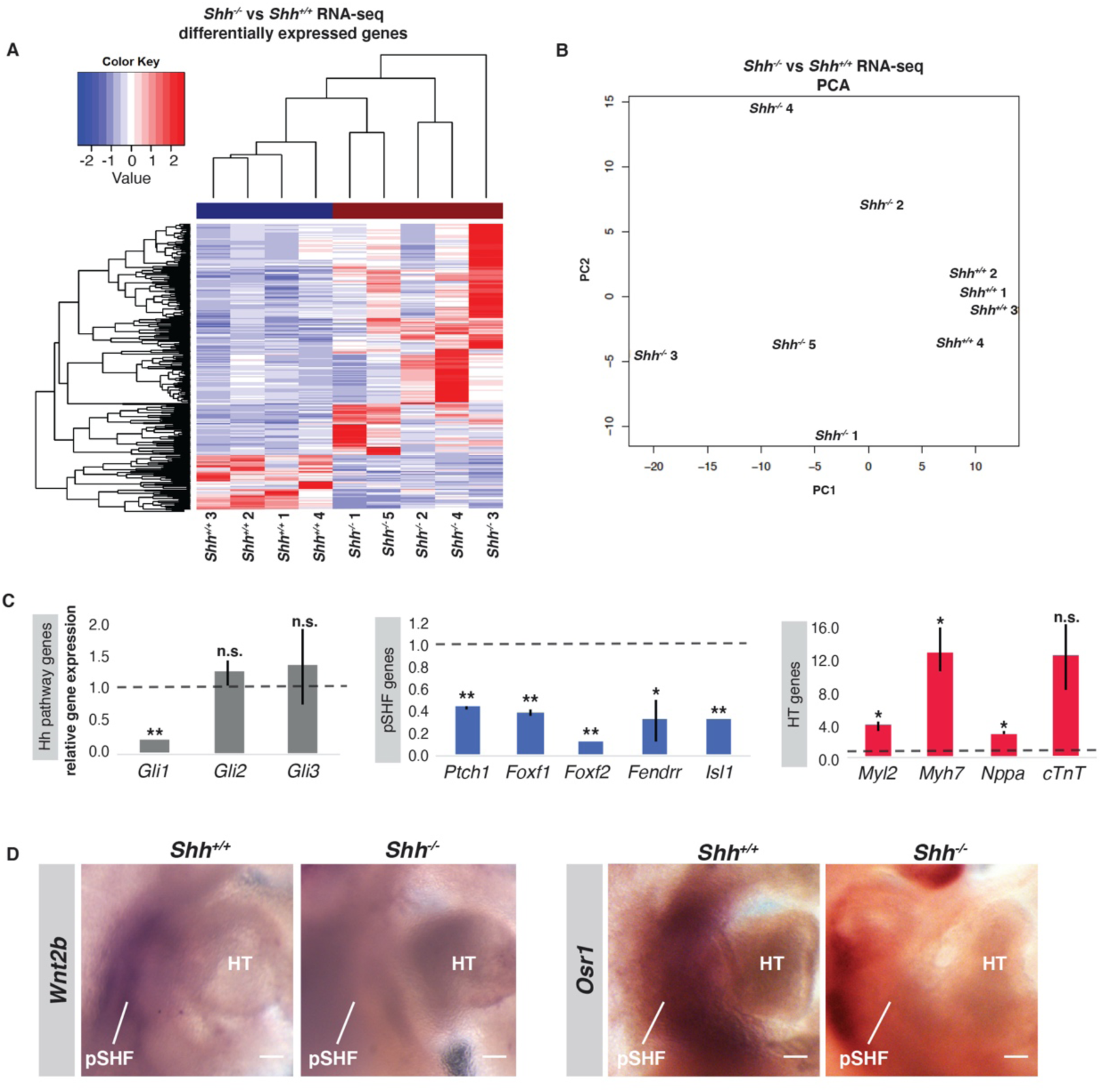
Differential gene expression analysis in the *Shh^-/-^* pSHF reveals disruption of the cardiac progenitor regulatory network. **(A)** Heatmap showing differentially expressed genes in *Shh^-/-^* replicate samples, relative to *Shh^+/+^* controls. **(B)** PCA plot of differentially expressed genes demonstrating that *Shh* genotype accounts for the highest proportion of variation among embryo gene expression profiles. **(C)** qPCR validation of reduced Hh pathway activation, reduced CP gene expression and increased CM gene expression in the *Shh^-/-^* pSHF (* *p*-value ≦ 0.05, ** *p*-value ≦ 0.01). **(D)** *In situ* hybridization demonstrating the expression domains of two pSHF marker genes (*Wnt2b* and *Osr1)* in the SHF of *Shh^+/+^* and *Shh^-/-^* embryos. Scale bar = 100μm. pSHF, posterior second heart field; HT, heart tube.

**Supplemental Figure 4.**
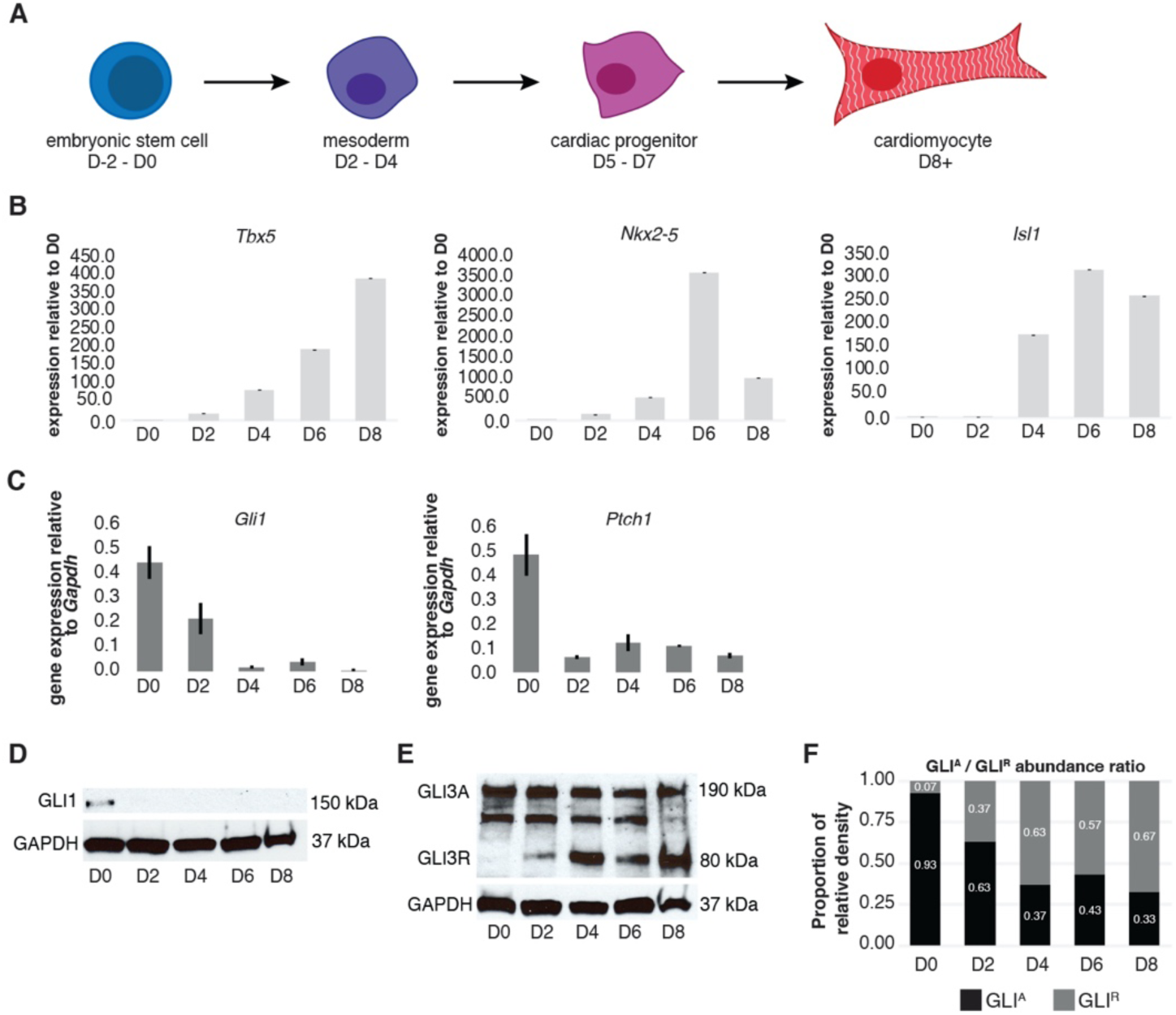
Hedgehog signaling is inactive in mESC-derived CPs and CMs. **(A)** Schematic depicting the *in vitro* mESC-derived CM differentiation system employed in these studies. **(B)** Relative expression levels of markers of the cardiac lineage during mESC-CM differentiation. **(C)** Relative expression levels of markers of active Hh signaling during mESC-CM differentiation. **(D)** Western blot demonstrating the expression level of GLI1 protein during mESC-CM differentiation. **(E)** Western blot demonstrating the expression level of GLI3^A^ (∼190 kDa) and GLI3^R^ (∼80 kDa) isoforms during mESC-CM differentiation. **(F)** Relative proportions of GLI^A^ and GLI^R^ proteins expressed during mESC-CM differentiation, based on western blot pixel density analysis.

**Supplemental Figure 5.**
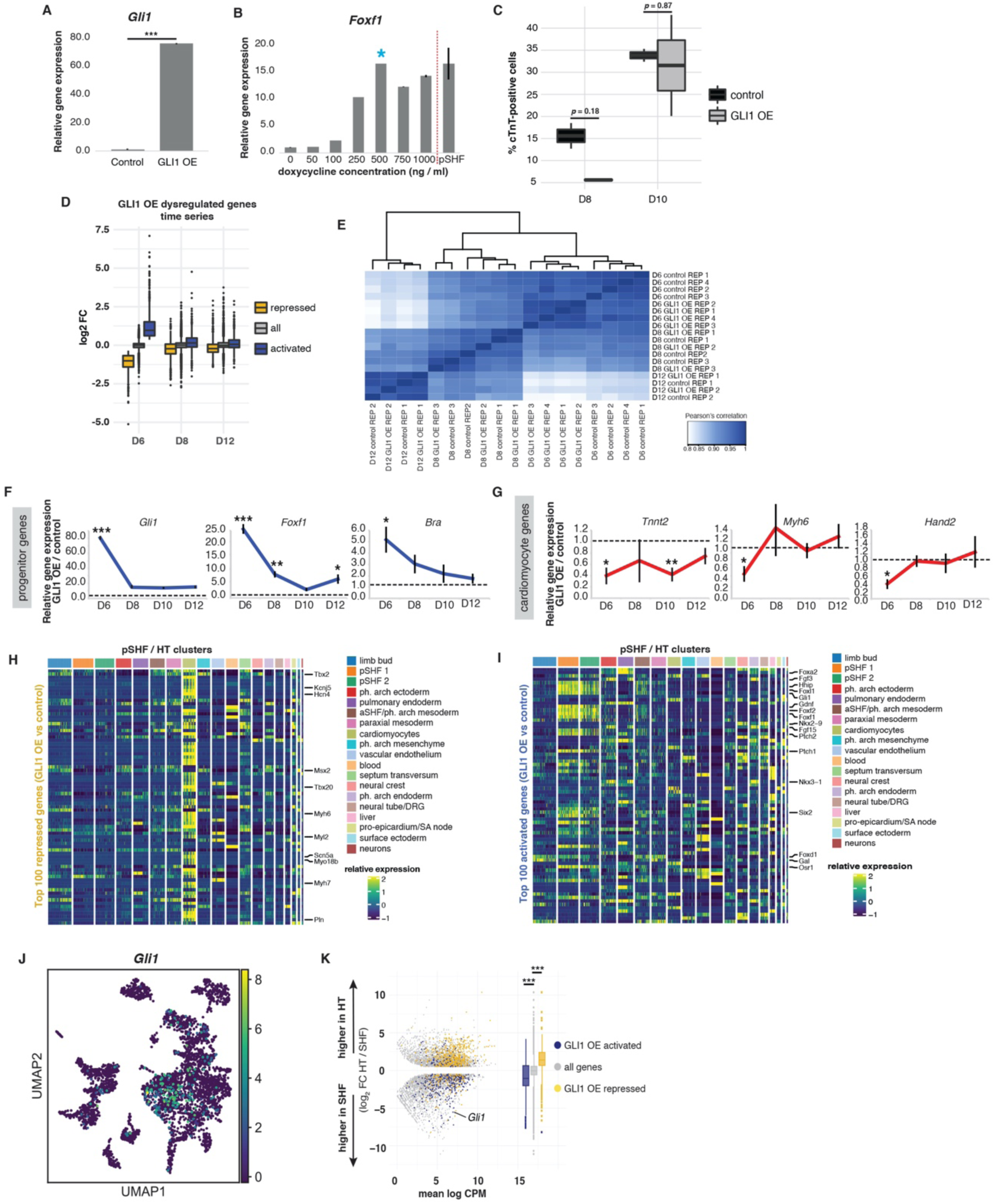
GLI1 OE in CPs leads to the transient activation of pSHF Hedgehog signaling targets but does not alter cell fate. **(A)** Relative gene expression analysis demonstrating that doxycycline treatment (dox, 500ng/ml) of the GLI1-FTA transgenic mESC line for 24 hours (D5-6) results in robust activation of *Gli1* transcript. **(B)** Expression levels of the Hh target *Foxf1*, relative to *Gapdh*, resulting from various dox concentrations in CPs, and as compared to endogenous *Foxf1* and *Gapdh* expression levels in the embryonic pSHF. **(C)** Percentage of cTnT+ cells in control and GLI1 OE samples at D8 and D10 of differentiation, as measured by fluorescence-activated cell sorting. **(D)** Time series of log_2_ fold changes of genes repressed and activated by GLI1 OE at D6, and their log_2_ fold changes at D8 and D12, relative to all expressed genes. The mean log2 fold changes of repressed, activated and all expressed genes were compared with a non-parametric Kruskall-Wallis rank sum test for D6, D8 and D12 samples: D6 χ_2_ = 3804.9, df = 2, *p*-value < 2e^−16^; D8 χ_2_ = 397.45, df = 2, *p*-value < 2e^−16^; D12 χ_2_ = 363.78, df = 2, *p*-value < 2e^−16^. **(E)** Heatmap displaying the Pearson’s correlation of gene expression profiles from GLI1 OE and control cells at D6, D8 and D12. **(F)** qPCR validation of the temporary activation of CP-specific genes in GLI1 OE cells relative to control cells (* *p*-value ≦ 0.05, ** *p*-value ≦ 0.01, *** *p*-value ≦ 0.005). **(G)** qPCR validation of the temporary repression of CM-specific genes in GLI1 OE cells relative to control cells (* *p*-value ≦ 0.05, ** *p*-value ≦ 0.01). **(H)** Heatmap of the wild-type E10.5 relative expression levels of the top 100 GLI1 OE CP repressed genes within 19 cell type clusters identified by Drop-seq. **(I)** Heatmap of the wild-type E10.5 relative expression levels of the top 100 GLI1 OE CP activated genes within 19 cell type clusters identified by Drop-seq. **(J)** UMAP plot 19 cell type clusters identified by Drop-seq indicating the expression of *Gli1* predominantly in the pSHF clusters. **(K)** MA plot and box plots illustrating the distribution of GLI1 OE dysregulated genes in the context of differential expression between the wild type E10.5 pSHF and HT (pairwise T-tests: activated vs all expressed genes *p-*value = 7.16e^−20^, repressed vs all expressed genes *p*-value = 1.49e^−57^).

**Supplemental Figure 6.**
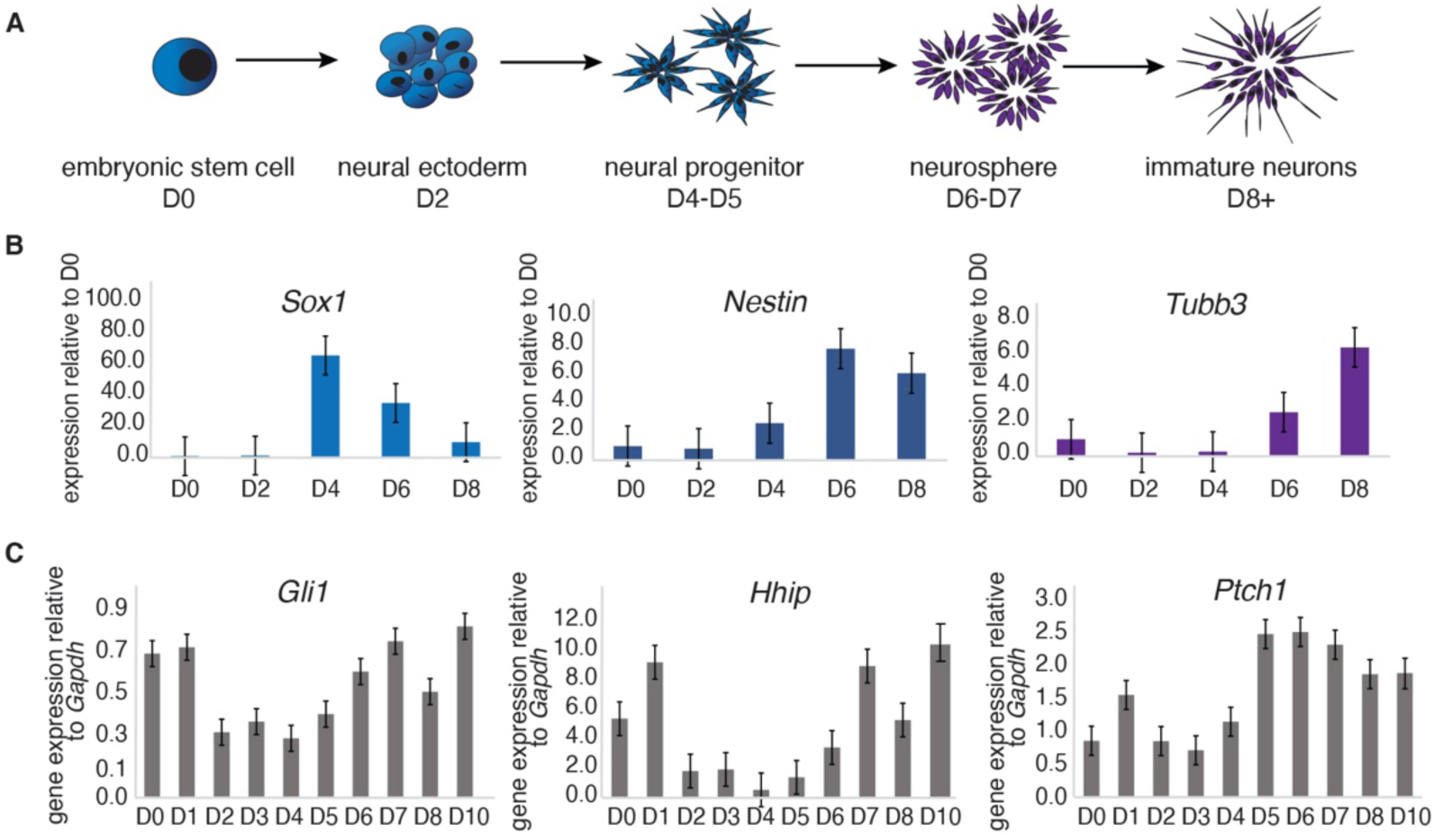
Hedgehog signaling is low in mESC-derived neural progenitors. **(A)** Schematic depicting the *in vitro* mESC-derived neural differentiation system employed in these studies. **(B)** Relative expression levels of markers of the neural lineage during neural differentiation. **(C)** Relative expression levels of markers of active Hh signaling during neural differentiation.

**Supplemental Figure 7.**
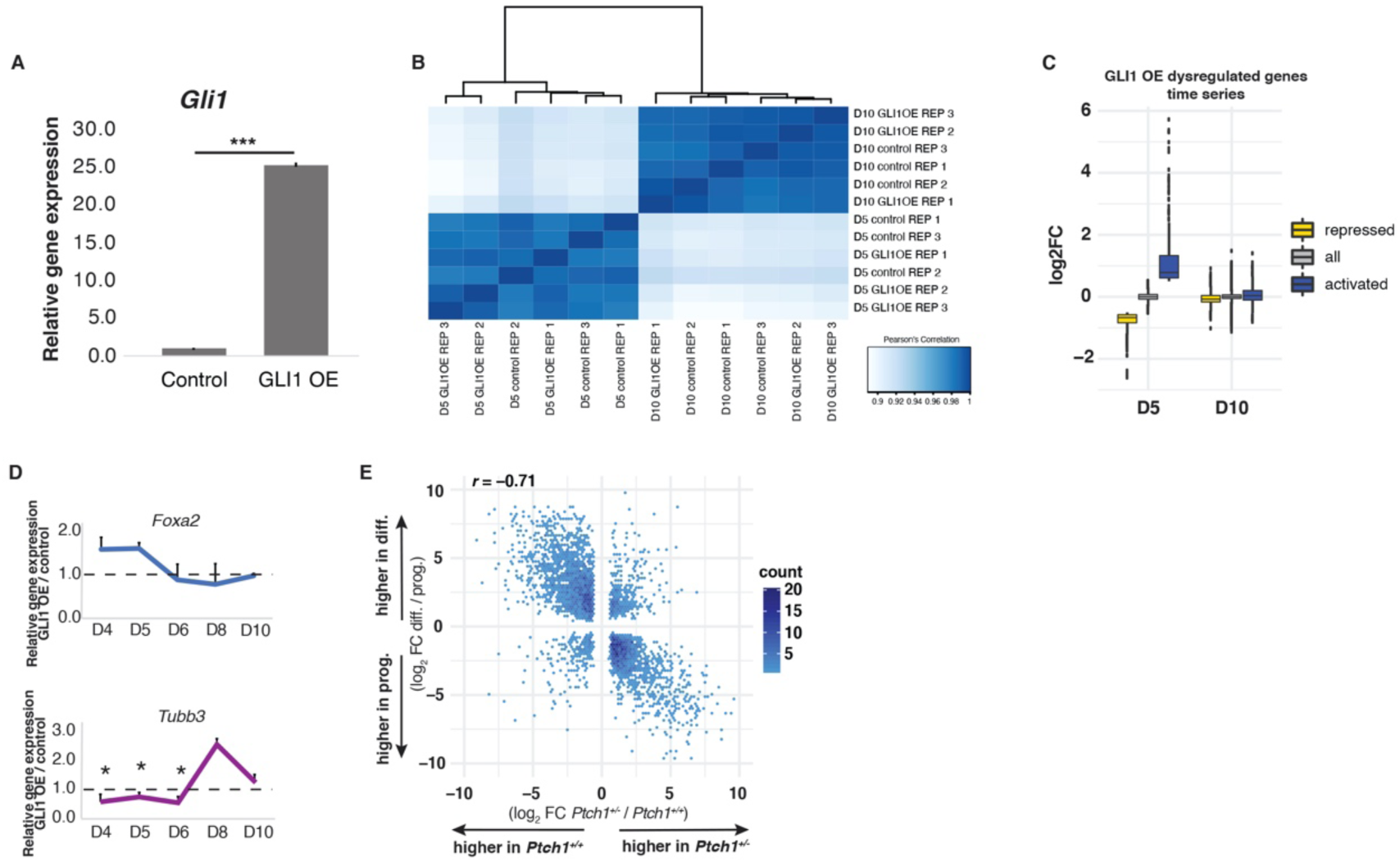
GLI1 OE in neural progenitors transiently activates neural progenitor gene expression, including FOX TFs. **(A)** Relative gene expression analysis demonstrating that dox treatment (500ng/ml) of the GLI1-FTA transgenic mESC line for 48 hours (D3-5) results in robust activation of *Gli1* transcript. **(B)** Heatmap displaying the Pearson’s correlation of gene expression profiles from GLI1 OE and control cells at D5 and D10. **(C)** Time series of log_2_ fold changes of genes repressed and activated by GLI1 OE at D5 and their log_2_ fold changes at D10, relative to all expressed genes. **(D)** qPCR validation of the temporary activation of a neural progenitor-specific gene (*FoxA2*) and repression of a differentiated neuron-specific gene (*Tubb3*) in GLI1 OE cells relative to control cells (* *p*-value ≦ 0.05). **(E)** Correlation plot comparing *Ptch1^+/-^* dysregulated fold changes to the fold changes of genes differentially expressed between ectodermal progenitors and differentiated neuron stages. Pearson correlation *(p-*value = < 2e^−16^).

**Supplemental Figure 8.**
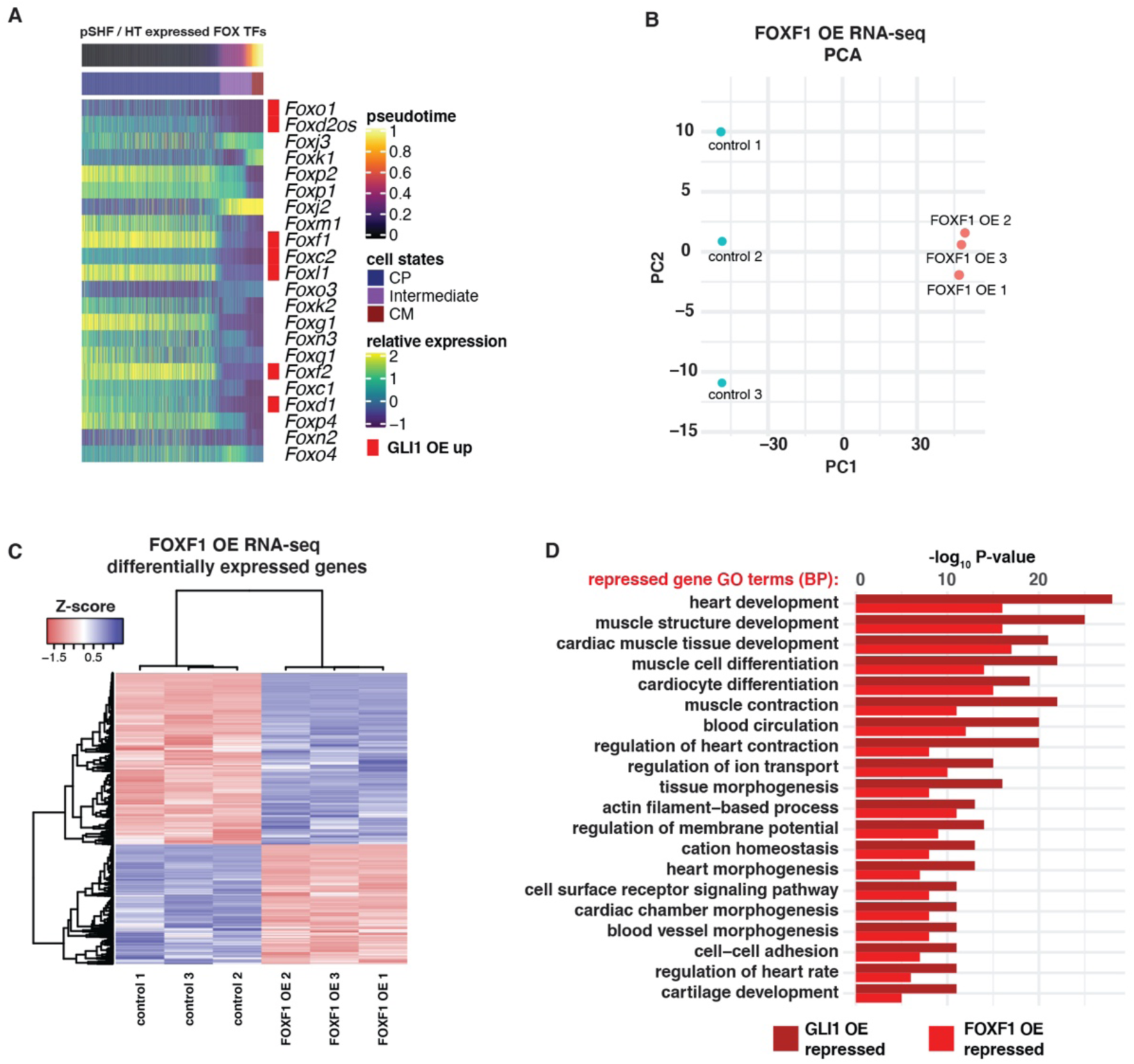
FOXF1 OE in CPs represses a similar cardiac differentiation program as GLI1 OE. **(A)** Heatmap showing denoised data from individual cells from cardiac-associated Drop-seq clusters indicating the expression levels of all FOX TFs expressed along the pseudotime differentiation trajectory. Cells are ordered on the x-axis by pseudotime and FOX TFs activated by GLI1 OE are indicated in red. CP, cardiac progenitor; CM, cardiomyocyte. **(B)** PCA plot of differentially expressed genes demonstrating that FOXF1 OE accounts for the highest proportion of variation among embryo gene expression profiles. **(C)** Heatmap showing differentially expressed genes in FOXF1 OE replicate samples, relative to control controls. **(D)** Comparative gene ontology (GO) analysis of GLI1 OE and FOXF1 OE repressed genes.

**Supplemental Figure 9.**
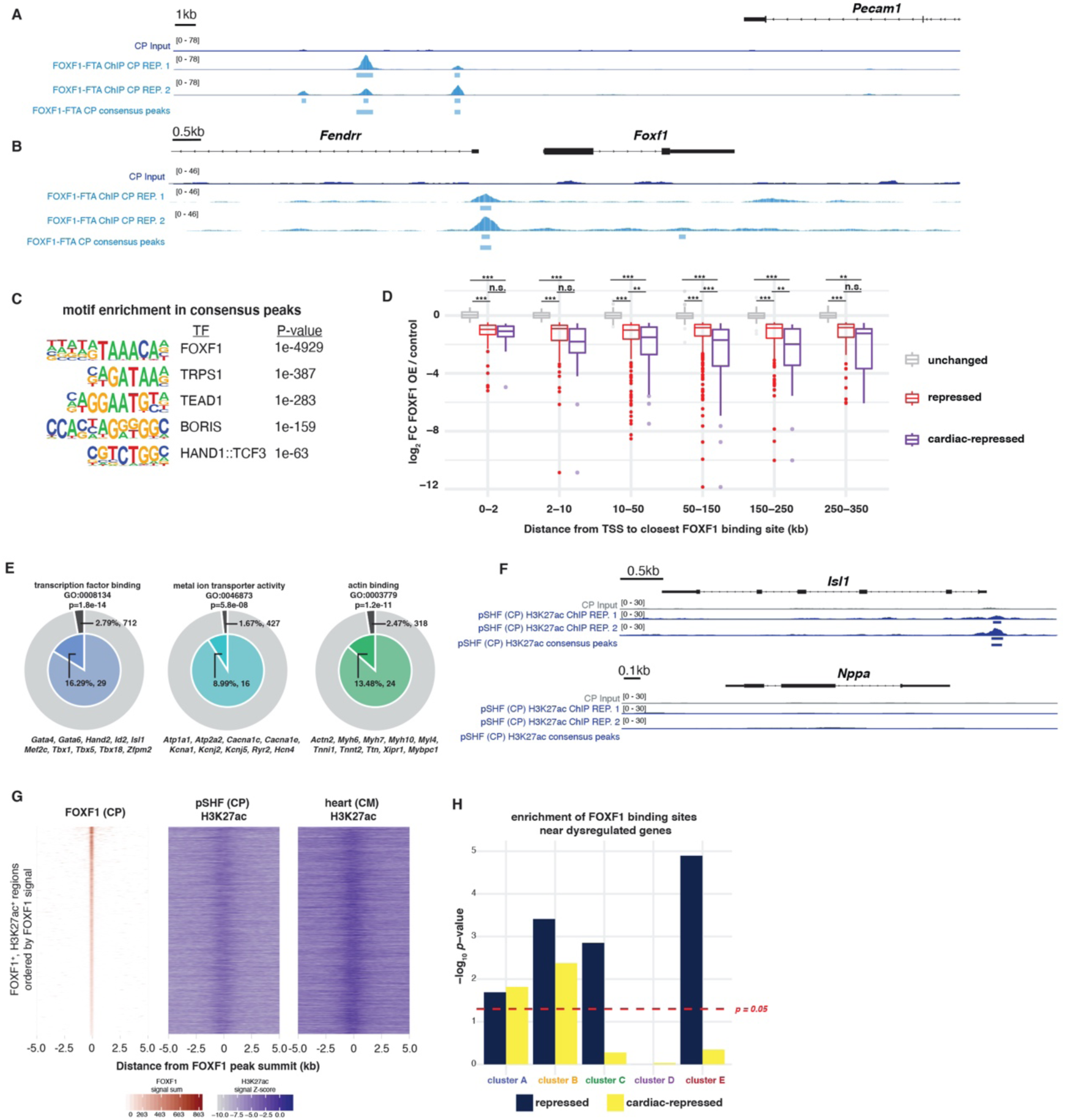
FOXF1 binds to enhancers in CPs that become activated in CMs. **(A)** Genome browser view of FOXF1-bound region near the previously-identified FOXF1 target, *Pecam1*, in replicate samples. **(B)** Genome browser view of FOXF1-bound region near the *Foxf1* and *Fendrr* promoters. **(C)** Binding motif logos of the top 5 most-enriched TF motifs found in sequences surrounding FOXF1-bound regions (summit +/− 1kb). **(D)** Boxplots showing the log_2_ FC of FOXF1 OE unchanged, all repressed and cardiac-repressed genes with various TSS distances from the nearest FOXF1 binding site (Kruskal-Wallis rank sum test, χ_2_ = 206.13 for 0-2 kb, χ_2_ = 278.93 2-10 kb, χ_2_ = 1025.6 10-50 kb, χ_2_ = 1233.3 50-150 kb, χ_2_ = 513.96 150-250 kb, χ_2_ = 243.32 250-350 kb, df = 2 for all tests, *p*-value = 2e^−16^ for all tests, followed by pairwise T-test with Benjamini-Hochberg correction, ** *p*-value ≦ 0.01, *** *p*-value ≦ 0.001). **(E)** Nested pie-charts demonstrating enrichment of genes associated with TF binding, metal ion transporter activity and actin binding molecular functions within FOXF1 OE repressed genes associated with FOXF1-bound regions within 250 kb. **(F)** Genome browser views of an H3K27ac-enriched region near a CP-specific gene and a lack of H3K27ac enrichment near a CM-specific gene, in replicate samples. **(G)** Heatmaps showing FOXF1 binding and H3K27ac signal z-scores in CPs and CMs for all FOXF1-bound cardiac enhancers. **(H)** Bar plot showing the enrichment of cardiac-repressed genes near FOXF1-bound regions in clusters A and B (one-sided Fisher’s Exact Test, dashed red line represents a *p*-value of 0.05).

## SUPPLEMENTAL TABLES

Supplemental Table 1: Top 100 differentially expressed genes in original 19 original cell-type clusters derived from E10.5 SHF/HT Drop-seq

Supplemental Table 2: Top 100 differentially expressed genes in the subset of 4 cardiomyocyte-associated clusters derived from E10.5 SHF/HT Drop-seq

Supplemental Table 3: 24 genes highly expressed in cardiomyocytes

Supplemental Table 4: Genes differentially expressed between combined CP cluster and CM cluster (log_2_ FC > 2.2, *p*-value < 0.05)

Supplemental Table 5: Genes differentially expressed between E10.5 Shh WT pSHF and E10.5 Shh KO pSHF (log_2_ FC > 0.5, FDR < 0.05)

Supplemental Table 6: Genes differentially expressed between E10.0 HT and pSHF (log_2_ FC > 0.5, FDR < 0.05)

Supplemental Table 7: Genes differentially expressed between GLI1 OE and control mESC-cardiac cells at D6, D8 and D12 (log_2_ FC > 0.5, FDR < 0.05)

7.1: D6 differentially expressed genes
7.2: D8 differentially expressed genes
7.3: D12 differentially expressed genes

Supplemental Table 8: Genes differentially expressed between GLI1 OE and control mESC-neural cells at D5 and D10 (log_2_ FC > 0.5, FDR < 0.05)

8.1: D5 differentially expressed genes
8.2: D10 differentially expressed genes

Supplemental Table 9: Genes differentially expressed between ectodermal progenitor tissue and differentiated neuronal tissue (log_2_ FC > 0.5, FDR < 0.05; data from ENCODE Project Consortium)

Supplemental Table 10: GO terms related to proliferation enriched with GLI1 OE-activated genes in neural progenitors

Supplemental Table 11: GO terms related to proliferation enriched with GLI1 OE-activated genes in pseudotime cardiac progenitor clusters

Supplemental Table 12: Genes differentially expressed between FOXF1 OE and control mESC-CPs at D6 (log_2_ FC > 0.5, FDR < 0.05)

Supplemental Table 13: Consensus regions bound by FOXF1 in mESC-CPs at D6 (present in both REP 1 + REP 2, mm10)

Supplemental Table 14: Data sets used in these studies from the ENCODE Project Consortium

Supplemental Table 15: GO term annotations used to identify cardiac-associated genes in these studies

## METHODS

### Animal Husbandry

Mouse experiments were completed according to a protocol reviewed and approved by the Institutional Animal Care and Use Committee of the University of Chicago, in compliance with the USA Public Health Service Policy on Humane Care and Use of Laboratory Animals. CD1 mice were obtained from Charles River Laboratories. The *Shh^-/-^* mouse line was obtained from the Jackson laboratory. *Osr1^eGFPCre-ERT2^* (Mugford et al., 2008), *Gli1^Cre-ERT2/+^* (Ahn and Joyner, 2004), and *ROSA26^Gli3R-Flag^* (Vokes et al., 2008, also called *ROSA26^Gli3T-Flag^*) lines were reported previously. *Foxf1^fl/fl^* (Ren et al., 2014) mice were obtained from the Kalinichenko lab (Cincinnati Children’s Hospital Medical Center).

### CD1 embryo Drop-seq analysis

SHF and HT tissue was microdissected from E10.5 CD1 embryos. Tissue from 40 embryos was pooled to generate Replicate 1 and tissue from 45 embryos was pooled to generate Replicate 2. Tissue was dissociated for 5 minutes at 37°C in TrypLE Express Enzyme reagent (Thermo Fisher 12604013), with shaking. The dissociation reagent was quenched with 10% FBS, and cells were subsequently washed in a 0.01% BSA/PBS solution. Two single cell RNA-seq libraries were generated using the standard Drop-seq protocol (Macosko et al., 2015) with minor modifications. Briefly, cells were filtered using a 40 µm mesh filter (Nalgene, USA) and suspended in 1x PBS and 0.01% BSA at a concentration 120,000/mL. Custom-synthesized Drop-seq barcode (BC) beads (Chemgenes, USA) at 100,000/mL were likewise filtered trough 100 µm mesh filter and suspended in Drop-seq Lysis Buffer. Using 3 mL syringes for each, the solutions were co-encapsulated in monodisperse droplets of 125 µm in an inert carrier oil (BioRad Evagreen) using microfluidic device fabricated in-house. The cells underwent lysis in the microfluidic droplets and the mRNA were captured onto the BC beads via polydT oligoes affixed to the BC beads. The droplets were then broken to retrieve the BC beads with mRNA hybridized onto them in high salt solution that stabilized the BC oligo+mRNA complexes. The beads are thoroughly washed and reverse transcription is performed to convert the BC+mRNA complex into barcoded cDNA. An Exonuclease I digestion is performed to digest any barcodes that are unbound to mRNA. PCR of 12-15 cycles is performed to amplify the barcoded single cell library, quantified and the quality of cDNA library is assessed. Sequencing libraries are prepared using Illumina Nextera kit with sample-specific indices added that allow de-multiplexing each sample from sequence data. The samples were sequenced on the Illumina NextSeq platform using a 75 cycle kit, with a custom Read1 primer (Macosko et al., 2015). The two biological replicates were processed on different days.

Raw sequencing data were processed through a bioinformatics pipeline based on the STARsolo method (Dobin et al., 2013) and as previously described (Selewa et al., 2020). Briefly, sequencing reads were aligned to the mouse reference genome (mm10) (Kent et al., 2002), and transcripts counts were quantified for each cell and gene. The pipeline is available here: https://github.com/aselewa/dropseqRunner. Data QC and analysis of the Drop-seq data was performed using the Scanpy toolkit (Wolf et al., 2018). Cells from both biological replicates were combined after removing cells with less than 1,100 detected genes and genes that were detected in less than 5 cells. We also excluded cells with more than 7,000 genes to avoid counting cell doublets and cells with more than 3% of reads mapped to mitochondrial genes indicative of low quality. This resulted in a final set of 3,824 high quality cellular transcriptomes. Read counts in each cell were normalized to a total of 10,000 reads and log transformed.

We used an iterative analysis strategy to first identify cardiac-associated cell types present in the entire dataset of cells from the SHF, and then repeated this process on a selected subset of cells corresponding to those with an atrial CM fate. First, we identified highly variable genes within the cell populations using default parameters, followed by PCA. Prior to clustering and non-linear dimensionality reduction, neighborhood graph was constructed using *n_neighbors=15* and *n_pcs=35.* We calculated both UMAP and tSNE representations for different subsets of the data and chose the representation that made facilitated visually distinguishing the independently derived clusters. Graph-based clustering was performed using the Leiden algorithm (Traag et al., 2019). We identified marker genes for each of the 19 original clusters and used them to assign cell types (Supplemental Table 1; Supplemental Figure 1D-E). All cardiac related cell types were then selected for further analysis to ensure that no CM cells were lost (clusters: pSHF1, pSHF2, aSHF/pharyngeal arch mesoderm, cardiomyocytes, septum transversum, pro-epicardium/SA node), yielding a subset of 1,402 cells (Supplemental Figure 1D-F and Supplemental Table 1). The entire analysis process was then repeated for this subset leading to 7 clusters representing cells in several cardiac lineages, which we further filtered to retain only cells definitively corresponding to the atrial CM lineage (*Gata4^+^, Nkx2-5^+^, Tbx5^+^, Tnnt2^+^*) (Han et al., 2019; de Soysa et al., 2019). This step excluded cells corresponding to the aSHF (*Tbx1^+^, Fgf10^+^*), the septum transversum (*Wt1^+^*) and the pro-epicardium/SA node (*Tbx18^+^*) (Supplemental Figure 1G-H). This final atrial CM-associated subset contained 846 cells sorted into 4 distinct clusters (Figure 1C-D and Supplemental Table 2). We confirmed the left/right side-specificities of the two progenitors (CP1 and CP2) by examining the expression of *D030025E07Rik* (right side marker) and *Pitx2* (left side marker) in the wild type pSHF *in vivo* (Supplemental Figure 2C-D).

To calculate the differentiation pseudotime we first computed a diffusion map using default settings (Haghverdi et al., 2016). To identify a starting cell for the pseudotime calculation, we calculated a ‘differentiation score’ using know cardiac marker genes (Supplemental Table 3) (de Soysa et al., 2019) and chose as the root cell the most differentiated cell according to this metric. We thus use 1-pseudotime to describe the pseudotime associated with the developmental progression. We chose this approach to overcome the larger heterogeneity within the progenitor population, which made choosing a progenitor root cell more difficult. Cell cycle scores were calculated as previously described (Tirosh et al., 2016) after converting the human gene symbols to the corresponding symbols for the mouse gene list. To confirm the identities of CPs and CMs, we identified differentially expressed genes based on a t-test using default settings in scanpy (Supplemental Table 4). To map activity of developmental signaling pathways, we obtained sets of target genes from 6 developmental pathways (Han et al., 2019) and used the aggregate expression in each cell as a proxy for pathway activity.

To display and cluster the expression patterns of these genes at the single cell level, we denoised the single cell expression measurements using the computational algorithm ENHANCE that relies on principal component analysis in order to separate biological variation from technical noise (Wagner et al., 2019). Denoised single cell gene expression values were displayed with heatmaps using the R package ComplexHeatmaps (Gu et al., 2016). Analysis for enriched GO terms was performed using Metascape (metascape.org, Zhou et al., 2019).

### *Shh*^-/-^ embryo transcriptome profiling

The pSHF was microdissected from five individual E10.5 *Shh^+/+^* and five *Shh^-/-^* embryos for bulk RNA-seq analysis, and yolk sacs were collected for genotyping. Tissues were mechanically homogenized in TRIzol Reagent (ThermoFisher Scientific 15596026), and RNA was isolated using RNeasy Mini RNA Isolation Kit (Qiagen 74104). 1ug of total RNA was then used to generate sequencing libraries using the TruSeq RNA Sample prep kit v2 (Illumina RS-122-2001), as per recommended instructions. Libraries were quantitated on an Agilent Bio-Analyzer and pooled in equimolar amounts. Pooled libraries were sequenced on the HiSeq2500 in Rapid Run Mode following the manufacturer’s protocols to generate stranded single-end reads of 50bp. The number of sequenced reads per sample ranged between 11.7 million 17.7 million with an average of 15 million sequenced per sample. Quality control for raw reads involved trimming the first 13bp with FastQ Groomer to ensure a median quality score of 36 or above for each sample. Fastq files were aligned to the UCSC mouse genome (mm9) (Kent et al., 2002) using TopHat (Cole Trapnell, 2009) (version 2.0.10) with the following parameters: (--segment-length 19 --segment-mismatches 2 --no-coverage-search). Between 11.4 million 17.2 million successfully mapped reads were then merged. One *Shh^+/+^* sample was discarded due to discordance with all other samples. Remaining samples were then analyzed for differential gene expression using Cuffdiff (Trapnell et al., 2013) (version 2.1.1) with quartile normalization. Significantly differentially expressed genes were identified using thresholds of FDR <0.05 and fold change > 1.5, resulting in 204 activated and 52 repressed genes, in the *Shh^-/-^* pSHF samples relative to *Shh^+/+^* samples. Differential expression of selected genes identified by RNA-seq was validated using qPCR. RNA was harvested from *Shh^+/+^* and *Shh^-/-^* pSHF samples with a Nucleospin RNA extraction kit (Machery-Nagel 740955.250). RNA was then used to perform one-step qPCR with the iTaq One-Step system (Bio-Rad 1725151), and expression levels in mutant samples were normalized to *Shh^+/+^* samples. *In situ* hybridization using RNA probes complimentary to *Wnt2b* and *Osr1* transcripts was performed as previously described (Hoffmann et al., 2014). GO term analysis was performed as described above. To intersect *Shh*-dependent genes with differentiation stage-dependent gene expression, we also performed bulk RNA-seq analysis on wild type embryonic tissues. The pSHF and HT from six CD1 embryos was microdissected and RNA-seq analysis was performed similarly to identify genes differentially expressed between the two tissues (fold change > 1.5, FDR < 0.05). Additional data analysis and visualization was performed in R.

### Gli1-FTA and FOXF1-FTA mESC line generation

To generate an inducible GLI1 mouse embryonic stem cell (mESC) line, the coding sequence for mouse *Gli1* or *Foxf1* was inserted in-frame with a Flag-Tev-Avi (FTA) tag into the *Hprt* locus of A2Lox.cre mES cells using a method previously described (Iacovino et al., 2011). Individual clones were assessed for differentiation potential and doxycycline-inducibility. One clone was selected and used for all experiments described herein, and all experiments were performed at the same passage number.

### mESC cardiomyocyte differentiation culture and GLI1 OE

mESCs were maintained and differentiated into CMs as previously described (Kattman et al., 2011). RNA was collected and qPCR was performed as above to assess the expression of CP markers *Tbx5, Nkx2-5* and *Isl1*, as well as Hh markers *Gli1*, and *Ptch1* relative to *Gapdh* in differentiating CMs. For western blots, protein was harvested with cell lysis buffer (Cell Signaling 9803), and 20-40 ug was loaded onto 4-15% pre-cast gels (Bio-Rad 17000546) and run using the Bio-Rad Mini-PROTEAN Tetra Cell system. Blots were probed with antibodies recognizing GLI1 (1:1,000; Cell Signaling 2534S), GLI3 (1:1,000; R&D AF3690) and GAPDH (1:10,000; Abcam ab8245). Pixel density for GLI1/3 was then computed, relative to the pixel density of GAPDH, using ImageJ (Schneider et al., 2012). The relative densities of GLI1/3 at various mESC-CM differentiation stages were summed, and the proportion of density contributed at each stage by GLI1 (GLI^A^) and GLI3 (GLI^R^) was calculated.

For GLI1 overexpression (GLI1 OE) experiments, doxycycline (Sigma D9891) was added to cultures at the CP stage (Day (D) 5). Cells were then washed and media was changed after 24hrs of exposure to doxycycline. qPCR and western blots were performed, as described above, on D6 samples to assess the expression level of *Gli1* and Hh targets, relative to untreated controls. Western blots for GLI1-FTA were probed with antibodies recognizing the FLAG epitope (1:1,000, F3165, M2) and GAPDH (as above). Expression of the Hh target, *Foxf1*, was quantified relative to *Gapdh* expression in cells treated with increasing concentrations of doxycycline and relative to *Gapdh* in pSHF embryo samples. Based on results from these assays, a final doxycycline concentration of 500ng/ml was chosen for all GLI1 OE experiments. mESC immunofluorescence was performed on control and GLI1 OE CMs with an antibody recognizing cardiac Troponin T (cTnT, 1:100, ThermoFisher Scientific MS-295-P1). The area of cTnT positivity was calculated as the mean grey area / mm^2^ of 5 fields of view across two biological replicates per condition, using the threshold measurement tool in ImageJ. The percentage of cTnT+ cells was calculated by dissociating D8 or D10 CMs and staining with a BV421-conjugated anti-cTnT antibody (BD Biosciences 565618, Lot 8136651, 13-11). The fluorescent signal from GLI1 OE and control cell samples stained for cTnT-positivity was normalized to fluorescent signal from cells stained with a BV421-conjugated isotype control IgG antibody (BD Biosciences 562438, Lot 8242926, X40). Fluorescence was quantified, and cells were counted on a BD LSRFortessa 4-15 HTS FACS instrument at the University of Chicago’s Cytometry and Antibody Technology core facility. The number of beating foci was calculated as the average number of independently beating regions within 5 fields of view across two biological samples, counted from 5 second videos taken on a Zeiss Axiovert 200m inverted widefield microscope at the University of Chicago’s Integrated Light Microscopy core facility.

RNA-seq was performed at D6, D8 and D12 of mESC-CM differentiations with at least two replicate GLI1 OE and control samples per stage. Reads were mapped to the mouse genome (mm9) (Kent et al., 2002) with Bowtie2 (Langmead and Salzberg, 2012) and transcripts were assigned and quantified with Cufflinks with default parameters (Trapnell et al., 2010). Reads were then normalized and differentially expressed genes for each timepoint were identified with edgeR (Mark D. Robinson, 2010). Differentially expressed genes were then filtered to include only genes with a log_2_ fold change ≥ 0.5 and an FDR ≤ 0.05. GO term analysis on D6 dysregulated genes, and qPCR validation of selected genes, was performed as described above. TF family enrichment was determined using TF family-membership annotations compiled in the Animal Transcription Factor Database (TFDB 3.0, http://bioinfo.life.hust.edu.cn/AnimalTFDB/#!/, Hu et al., 2019).

### mESC neuronal differentiation culture and GLI1 OE

Neuron differentiation was performed as previously described (Sasai et al., 2014; Ying and Smith, 2003). Briefly, cells were plated at a density of 9.5 × 10^4^ /cm^2^ on a 0.1% gelatin-coated dish and allowed to differentiate in N2B27 medium, which was replaced every two days. RNA was collected and qPCR was performed as described above to assess the expression of neural stem cell marker *Sox1*, neural progenitor markers *Nestin* and *Pax6* and neuron marker *Tubb3,* as well as Hh markers *Gli1*, *Hhip* and *Ptch1* relative to *Gapdh* in differentiating neurons.

For GLI1 overexpression experiments, 500ng/ml doxycycline (Sigma D9891) was added to cultures at the neural progenitor stage (Day 3). Cells were then washed and media was changed after 48hrs of exposure to doxycycline. The number of axons and the number of neurospheres / neural rosette clusters were manually counted by two independent observers blinded to the treatment from ten fields of view across two biological replicates using 10x brightfield microscopy (Olympus IX81). mESC immunofluorescence was performed on control and GLI1 OE cells with an antibody recognizing β-Tubulin III (*Tubb3*/TUJ1, 1:100, R & D Systems MAB1195). The area of TUJ1 positivity was calculated as the mean grey area / mm^2^ of 5 fields of view across two biological replicates per condition, using the threshold measurement tool in ImageJ.

RNA-seq was performed with mESC neural GLI1 OE and control samples at D5 and D10 in a similar manner as for mESC cardiac samples. qPCR validation of dysregulated genes was performed, as above, as were GO term analysis and TF enrichment analysis. A circos plot to intersect cardiac and neural GLI1-activated genes was generated using Metascape. Additional comparative analyses and data visualization was performed in R.

To compare genes dysregulated in the *Ptch1^+/-^* mouse medulla to stage-dependent ectodermal differentiation genes, we first obtained ectodermal progenitor and differentiated neuron samples from the ENCODE Project Consortium (Supplemental Table 14) (ENCODE Project Consortium, 2012) and performed differential expression testing with DESeq2 (version 1.20) and parameter minReplicatesForReplace set to FALSE to disallow the software from omitting samples. We then compared the stage-dependent changes in gene expression (log_2_ fold change progenitor / differentiated) to the reported log_2_ fold change of genes differentially expressed in the *Ptch1^+/-^* mouse medulla, relative to *Ptch1^+/+^* samples (Grausam et al., 2017). Correlation was tested using R’s cor.test function with the Pearson method.

### mESC cardiomyocyte differentiation culture and FOXF1 OE

Differentiation markers, and the effect of FOXF1 OE in CPs, was assessed in the FOXF1-FTA cell in the same manner as for the GLI1-FTA cell line, however, a doxycycline concentration of 250 ng/ml was used to induce FOXF1 expression. Cells were harvested after 24 hours of FOXF1 OE (D5-6) and RNA was harvested for RNA sequencing, as above for GLI1 OE. Analysis was performed as above, with some exceptions: reads were aligned with STAR v2.5.1 to mouse genome mm10 (Kent et al., 2002), version 19 from GENCODE. We used Samtools v1.5 to remove reads with phred quality score below 30. Reads were counted with HTSeq v0.6 in union mode with default settings. Differential expression test was performed with edgeR (3.30.3), and results were visualized with ggplot2 (v3.3.2). Genes that were associated with cardiac development or cardiac function were defined as dysregulated genes annotated to any of the GO terms listed in Supplemental Table 15.

### FOXF1 binding localization in CPs

The GLI1-FTA cell line was used to induce the expression of native FOXF1 protein to identify FOXF1 binding sites in CPs. Cells at the CP stage (D6) were treated with doxycycline (500 ng/ml) for 24 hours and harvested at D7. Approximately 2 x 10^6^ cells were cross-linked in 1% PFA for 5 minutes at room temperature, with rocking. The reaction was then quenched with 125 mM glycine. The cross-linked tissues were incubated in LB1 (50 mM HEPES-KOH, pH 7.5, 140 mM NaCl, 1 mM EDTA, 10% Glycerol, 0.5% NP-40, 0.25% Triton X-100) with protease and phosphatase inhibitors on ice for 10 minutes. The tissues were then sonicated in LB3 (10 mMTris-HCl, pH 8.0, 100 mM NaCl, 1 mM EDTA, 0.5 mM EGTA, 0.1% sodium deoxycholate, 0.5% N-lauroyl sarcosine, 1% Triton X-100) with protease and phosphatase inhibitors for 5 minutes at 4°C. Chromatin extract was then cleared by centrifugation at 14,000g, 4°C for 10 minutes. For immunoprecipitation, the chromatin extract was incubated with 10ug of an antibody recognizing FOXF1 (R&D Systems AF4798, Lot CAHH0316111) at 4°C for >12 hours in a total volume of 200 μL. The immune-complexes were captured by Protein G-conjugated magnetic beads (Life Technologies, 1003D), washed in sequence by LB3, LB3 with 1 M NaCl, LB3 with 0.5 M NaCl, LB3, and TE (10 mM Tris-HCl, pH 8.0, 1 mM EDTA). The captured chromatin, and input samples, were eluted in ChIP Elution Buffer (10 mM Tris-HCl, pH 8.0, 1 mM EDTA, 1% SDS, 250 mM NaCl) at 65°C. After RNase and proteinase K treatment and reverse cross-linking, DNA was purified.

High-throughput sequencing libraries from ChIP and input DNA were prepared using the NEBNext Ultra DNA Library Prep Kit (New England Biolabs, E7370S). During library preparation, adaptor-ligated DNA fragments of 200-650 bp in size were selected before PCR amplification using Sera-Mag magnetic beads (GE, 6515-2105-050-250). DNA libraries were sequenced using Illumina Hi-seq instruments (single-end 50 base) by the Genomics Core Facility at the University of Chicago. Reads were aligned to mouse genome mm10 (Kent et al., 2002) from UCSC with Bowtie2 v2.3.2. Reads with a phred score less than 30, aligned to mitochondrial genome and clone contigs were removed with Samtools v1.5. Reads were sorted with Samtools sort by genomic location before removing duplicates with picardtools MarkDuplicates v2.8.1 with default settings (http://broadinstitute.github.io/picard/). Peaks were called on uniquely mapped reads with macs2 (v2.1.1 settings -f BAM -g mm -B -q 0.05), grouping together both replicates as well as calling peaks on individual samples. Consensus peaks were determined by intersecting individual peak calls with Homer mergePeaks v4.1.1. Consensus peak summits were determined by intersecting the consensus peak regions with the summit file from the grouped peak calls. Coverage file (bigwig) was generated with macs2 (same version as above) using the bdgcmp function (-m FE) followed by bedGraphToBigWig v4 from UCSC. FOXF1 ChIP peaks were associated with FOXF1 OE dysregulated genes by identifying all dysregulated genes containing a TSS within 250kb of a FOXF1 ChIP peak region using the Bedtools makewindows tool (v2.29.1).

### Active enhancer identification in the embryonic pSHF

To identify regions enriched for H3K27ac histone modification, putatively representing active enhancer regulatory elements in CPs, we performed ChIP-seq on embryonic pSHF tissue. We immunoprecipitated H3K27ac+ DNA following the protocol described for FOXF1 ChIP-seq above, with two major exceptions: 1) chromatin was prepared from microdissected pSHF tissue from E9.5 CD-1 mouse embryos (2x from 50 embryo pools each), and tissues were cross-linked in PBS containing 1% formaldehyde at 25°C for 10 minutes and 2) the chromatin extract was incubated with 1ug of anti-H3K27ac antibody (Wako Chemicals 306-34849, MABI0309, Lot 14007) prior to capture, washing and elution. Library prep and sequencing was performed as described above.

Raw sequencing reads were aligned to the mm10 genome (Kent et al., 2002) using Bowtie2 and SAMtools (Heng Li, 2009) requiring a minimum mapping quality of 10 (-q 10). Pooled peak calling was performed using default settings of MACS2 callpeak (Feng et al., 2012; Zhang et al., 2008) with a q-value set to 0.05 and tag size set to 6 (-q 0.05 -s 6). A fold-enrichment track was generated using MACS2 using the bdgcmp function (-m FE) for visualization on the IGV genome browser (v2.4.14,) (Robinson et al., 2011). Peaks overlapping with ENCODE blacklist sites (ENCODE Project Consortium, 2012) were removed with the Bedtools intersect tool (Quinlan and Hall, 2010).

pSHF H3K27ac+ peak regions (summits +/− 1kb) were intersected with H3K27ac+ regions (summits +/− 1kb) from the adult mouse heart obtained from the ENCODE Consortium (Supplemental Table 14), and with our FOXF1-bound regions (summits +/− 1kb) in CPs to identify putative FOXF1-bound cardiac enhancers. Bedgraph files from all three ChIP data sets were used to quantify the signal within FOXF1-bound cardiac enhancers using the Bedtools map tool with the -o sum option. FOXF1-bound regions were clustered based on H3K27ac z-score normalized signal in R with k-means clustering to identify dynamic enhancer groups. The closest FOXF1 OE dysregulated gene to each region was identified with the Bedtools closest tool, and GO term analysis on the closest dysregulated genes for each category was performed with Metascape. IGV was used to visualize FOXF1 binding and H3K27ac ChIP enrichment at individual loci.

### Embryo Immunofluorescence & Histology

Embryos were harvested from timed pregnancies and yolk sacs were collected for genotyping. For *Osr1^eGFPCre-ERT2/+^*; *ROSA26^Gli3R-Flag/+^* (*Gli3^R^* OE) and *Gli1^Cre-ERT2/+^*; *Foxf1^fl/fl^* (*Foxf1* cKO) embryos, pregnant female mice were administered 2 mg tamoxifen at E8.5 and E9.5 (*Gli3^R^* OE) or E7.5 and E8.5 (*Foxf1* cKO) by intraperitoneal injection. Embryos were then harvested at E10.5 for immunofluorescence studies, or at E14.5 for histological examination. For immunofluorescence, E10.5 embryos were fixed overnight in 4% paraformaldehyde, washed with PBS and processed for paraffin sectioning. 5um serial sections were generated and used for immunofluorescence with primary antibodies recognizing sarcomeric myosin (1:20, DSHB MF20) and Alexa Fluor-conjugated secondary antibodies (1:1000, Thermo Fisher Scientific). Antigen retrieval was performed on all sections with 10mM sodium citrate buffer. DAPI was used to counterstain tissues and provide a tissue reference. Sections were imaged with an Olympus IX81 inverted widefield microscope using 10x and 20x objectives in the University of Chicago’s Integrated Light Microscopy Core Facility. Images were processed with ImageJ.

Histological studies were performed at E14.5. Embryos were harvested and fixed in 4% paraformaldehyde overnight at 4°C, and then processed for paraffin sectioning. 5um sections were then stained with hematoxylin and eosin to reveal the structural morphology of the atrial septum in mutant and control embryos.

## DATA DEPOSITION

Bulk and single-cell RNA-sequencing and ChIP-sequencing data will be deposited in the Gene Expression Omnibus databank upon publication.

## AUTHOR CONTRIBUTIONS

M.R. designed and performed the experiments and wrote the manuscript. C.P-C. and S.P. also designed experiments. A.R., J.J-L, A.G., J.D.S., A.D.H., S.L., H.E., and M.K performed *in vivo* experiments. A. R., S.H., J.J-L., N.D., E.L., C.K., S.Y., and E.H. performed *in vitro* experiments. S.P., C.P-C., X.H.Y., S.I. and K.I. performed computational analyses. S.S-K.C. and M.K. generated the GLI1-FTA and FOXF1-FTA mESC lines. D.J.G., S.P. and A.B. designed the experiments. I.P.M. designed the experiments and wrote the manuscript.

## ACKNOWLEDGEMENTS

The authors are deeply grateful to the staff of the Genomics, DNA Sequencing, Integrated Light Microscopy, Cytometry and Antibody Technology and Human Tissue Resource Center core facilities at the University of Chicago. This work was supported by the NIH/NHLBI: R01 HL092153 (I.P.M), R01 HL124836 (I.P.M), R01 HL126509-02 (I.P.M), R01 HL147571-01 (I.P.M), NIH/NHLBI NRSA T32 HL007381-36/37 (M. R.) and NIH/NHLBI F32 HL136168-01 (M. R.). This work was also supported by the American Heart Association: AHA 13EIA14690081 (I.P.M.) and AHA Postdoctoral Fellowship 17POST33670937 (M. R.).

